# MAPK signaling modulates the partition of DCP1 between P-bodies and stress granules in plant cells

**DOI:** 10.1101/2024.10.31.621288

**Authors:** Siou-Luan He, Ying Wang, Libo Shan, Ping He, Jyan-Chyun Jang

## Abstract

Processing bodies (PBs) and stress granules (SGs) are membrane-less cellular compartments consisting of ribonucleoprotein complexes. Whereas PBs are more ubiquitous, SGs are assembled mainly in response to stress. PBs and SGs are known to physically interact and molecules exchange between the two have been documented in mammals. However, the molecular mechanisms underpinning these processes are virtually unknown in plants. We have reported recently that tandem CCCH zinc finger 1 (TZF1) protein can recruit MAPK signaling components to SGs. Here we have found that TZF1-MPK3/6-MKK4/5 form a protein-protein interacting network in SGs. The mRNA decapping factor 1 (DCP1) is a core component of PBs. MAPK signaling mediated phosphorylation triggers a rapid reduction of DCP1 partition into PBs, concomitantly associated with an increase of DCP1 assembly into SGs. Furthermore, we have found that plant SG marker protein UBP1b (oligouridylate binding protein 1b) plays a role in maintaining DCP1 in PBs by suppressing the accumulation of MAPK signaling components. Together, we propose that MAPK signaling and UBP1b mediate the dynamics of PBs and SGs in plant cells.

## Introduction

Processing bodies (PBs) and stress granules (SGs) are two types of cytoplasmic biomolecular condensates that dynamically assembled in response to environmental stresses. Dysregulation of PBs and SGs has been implicated in various human diseases such as neuro- and muscular- degenerative diseases, neuro-developmental diseases, and cancers (Riggs *et al*, 2020; Ripin & Parker, 2023). Although previous studies have suggested that PBs and SGs perform distinct functions, as each contains a unique set of proteins, multiple observations indicate that SGs interact with PBs and likely exchange messenger ribonucleoproteins (mRNPs) between each other (Buchan & Parker, 2009). Under specific stress conditions, PBs often dock with SGs, and the overexpression of certain proteins that localize to both structures can lead to the fusion of PBs and SGs (Stoecklin & Kedersha, 2013). For example, the overexpression of tristetraprolin (TTP) or butyrate response factor-1 (BRF-1), two RNA-binding proteins that target ARE-containing mRNAs to PBs for degradation (Franks & Lykke-Andersen, 2007) or cytoplasmic polyadenylation element binding protein (CPEB1), another dual SG/PB protein (Wilczynska *et al*, 2005), lead to the tight clustering of PBs around and within SGs (Kedersha *et al*, 2005; Wilczynska *et al*., 2005). Overexpression of ubiquitin-associated protein 2-like (UBAP2L), and the interaction between UBAP2L and SG and PB nucleating protein such as G3BP (stress granule assembly factor) and DDX6 (DEAD-box RNA helicase), respectively, induces hybrid granules containing SG and PB components in the cells (Riggs *et al*, 2024). Furthermore, PBs can play a role in promoting SG assembly by providing untranslated mRNAs, shared proteins, and translational repressors that are necessary for nucleating SGs during cellular stress conditions (Buchan *et al*, 2008). On the other hand, SGs could still form in the absence of PBs. DDX6 (Rck/p54) is an evolutionarily conserved member of the DEAD-box RNA helicase family involved in the inhibition of translation and storage and the degradation of cellular mRNAs in PBs. When DDX6 expression is reduced, PB formation is strongly impaired, whereas DDX6 knockdown causes the PB-specific protein DCP1 to relocate to SGs (Serman *et al*, 2007), suggesting that some proteins involved in PB assembly may switch roles and contribute to co-aggregate with SG proteins under specific stress conditions.

In plants, RNA-binding protein 47b (Rbp47b) and Tudor Staphylococcal Nuclease (TSN2) are considered as core components of plant SGs (Maruri-Lopez *et al*, 2021). Rbp47b and TSN2 interactome analysis in Arabidopsis revealed the presence of mitogen-activated protein kinase (MAPK) signaling components MPK3, MKK4, and MKK5 (Solis-Miranda *et al*, 2023), suggesting that MAPK signaling could mediate SG dynamics, perhaps by phosphorylating key SG components. MAPK cascades (including MPK3 and MPK6 and their upstream regulators MKK4 and MKK5) play a crucial role in plant lifecycle, where they regulate a wide range of physiological processes, including growth and development as well as in responses to environmental cues such as cold, heat, drought, and especially pathogen attack (Zhang & Zhang, 2022). Some other kinases can also be recruited to SGs and PBs (Lopez-Palacios & Andersen, 2023; Shah *et al*, 2014). The SG component Ras-GAP SH3-binding protein (G3BP1) recruits casein kinase 2 (CK2) to SGs and phosphorylation of G3BP1 by CK2 promotes dissociation of G3BP from SGs and triggers SG disassembly (Reineke *et al*, 2017). The yeast kinase Sky1 is recruited to heat-induced SGs, where it can phosphorylate substrates Npl3 (a nucleocytoplasmic mRNA shuttling protein) to promote SG dissolution (Shattuck *et al*, 2019). SGs also play a role in sequestering signaling molecules as a protective mechanism. For example, high-heat stress stimulates MAPK activation, which causes fission yeast protein kinase C (Pck2) translocation from the plasma membrane into SGs to suppresses MAPK hyperactivation and cell death (Sugiura, 2021).

In plants, the molecular mechanisms underpinning the interaction and equilibrium between PBs and SGs are unclear. Here we present multiple lines of evidence to propose that MAPK signaling pathway is involved in triggering a major PB component DCP1 to relocate to SGs. We previously showed that tandem CCCH zinc finger 1 (TZF1) recruits MAPK signaling components to SGs (He *et al*, 2024). Using a high throughput and high-fidelity Arabidopsis protoplast transient expression system (He *et al*., 2024), we have found that MPK3, MPK6, MKK4, and MKK5 form homo- and hetero-dimers with each other in SGs. DCP1 is phosphorylated by MPK3 and MPK6 and the kinase activity of MKK5 is required for DCP1 to localize to SGs. In support of this notion, the phospho-dead form of DCP1^S237A^ is mainly localized in PBs and phosphor-mimetic form of DCP1^S237D^ is mainly localized in SGs. Oligouridylate binding protein 1b (UBP1b) is an RNA-binding protein and it has been used widely as an SG marker. Surprisingly, MAPK signaling-mediated reduction of DCP1 sequestration to PBs is antagonized by the SG marker UBP1b, as UBP1b suppresses the accumulation of MPK3/6 and MKK4/5. Consistent with this idea, the abundance of DCP1 PBs is enhanced by the co-expression of UBP1b, as well as an additional mechanism independent of DCP1’s phosphorylation status. Together, our results indicate that MAPK phosphorylation could serve as a switch for DCP1 sequestration from PBs to SGs. This pathway is counteracted by an SG marker UBP1b that could diminish the level of MAPK signaling components.

## Results

### TZF1-MPK3/6-MKK4/5 interacting network

We have shown previously that TZF1 could interact with MPK3/6 and MKK4/5 in stress granules (He *et al*., 2024). Here we have found that MPK3 and MPK6 could interact with itself and each other in bimolecular fluorescence complementation (BiFC) analysis. MKK4 and MKK5 could also interact with itself and each other (Fig. 1A). Furthermore, MPK3 or MPK6 could also interact with MKK4 or MKK5 in BiFC analysis (Fig. 1B). Remarkably, all the interacting BiFC signals were co-localized with SG localized TZF1 in cytoplasmic granules, suggesting that MPK3/6 and MKK4/5 interact in SGs. Interestingly, when the BiFC analysis was conducted in the presence of nuclear marker NLS-RFP, the interaction between MPK3 or MPK6 with MKK5 was mainly in the nucleus (Fig. 2D), as opposed to in the cytoplasmic granules when co-expressed with TZF1 (Fig. 1A-B). However, BiFC signals of MKK4 and MKK5 self- and cross-interactions remained distinctively in cytoplasmic granules (Fig. 2D). These results suggest that TZF1 recruits MPK3/6-MKK4/5 interactions to SGs. In fact, the BiFC signals of MPK3/6 and MKK5 cross-interactions were mostly localized in the nucleus when co-expressed with NLS-RFP (Fig. 2D) and TZF1 co-expression could similarly recruit these interactions to SGs (Fig. 1B).

**Figure 1.**
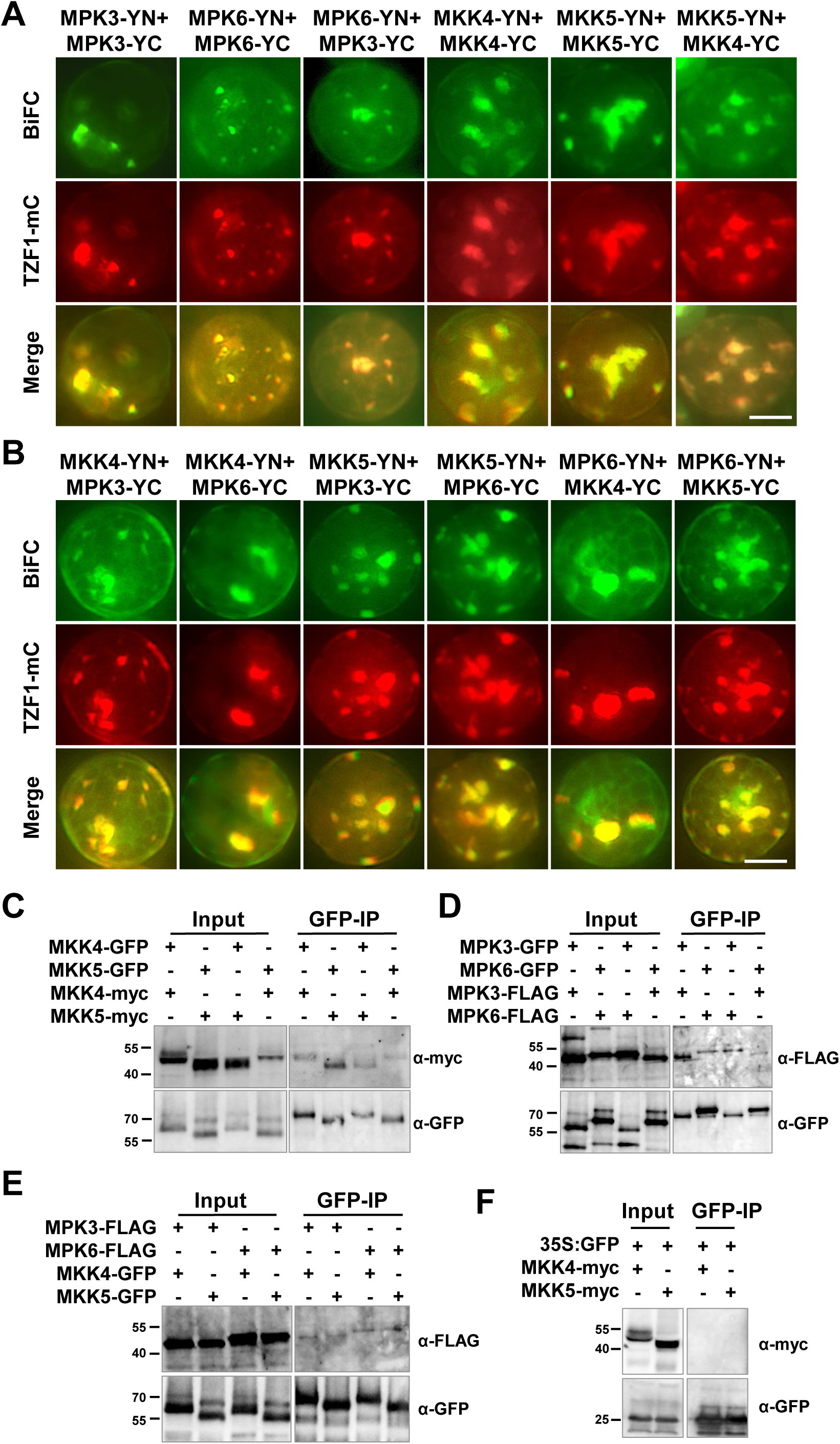
Protein-protein interaction of MPK3/6 and MKK4/5 in BiFC and Co-IP analyses. **(A)** Self- and cross-interaction of MPK3 with MPK6, and MKK4 with MKK5. The BiFC signals are completely co-localized with TZF1. **(B)** MKK4/5 and MPK3/6 cross-interact with each other. The BiFC signals are completely co-localized with TZF1. Scale bar= 10 μm. **(C-F)** Protein- protein interaction of MPK3/6 and MKK4/5 in Co-IP analyses. DNA constructs were co- expressed in an Arabidopsis protoplast transient expression analysis. GFP antibody was used for immunoprecipitation and immunoblot analyses were performed using various antibodies as indicated. **(C)** MKK4 and MKK5 self- and cross-interaction. **(D)** MPK3 and MPK6 self- and cross-interaction. **(E)** MKK4/5 and MPK3/6 cross-interaction. **(F)** Negative controls showing no interaction between MKK4/5 and GFP.

**Figure 2.**
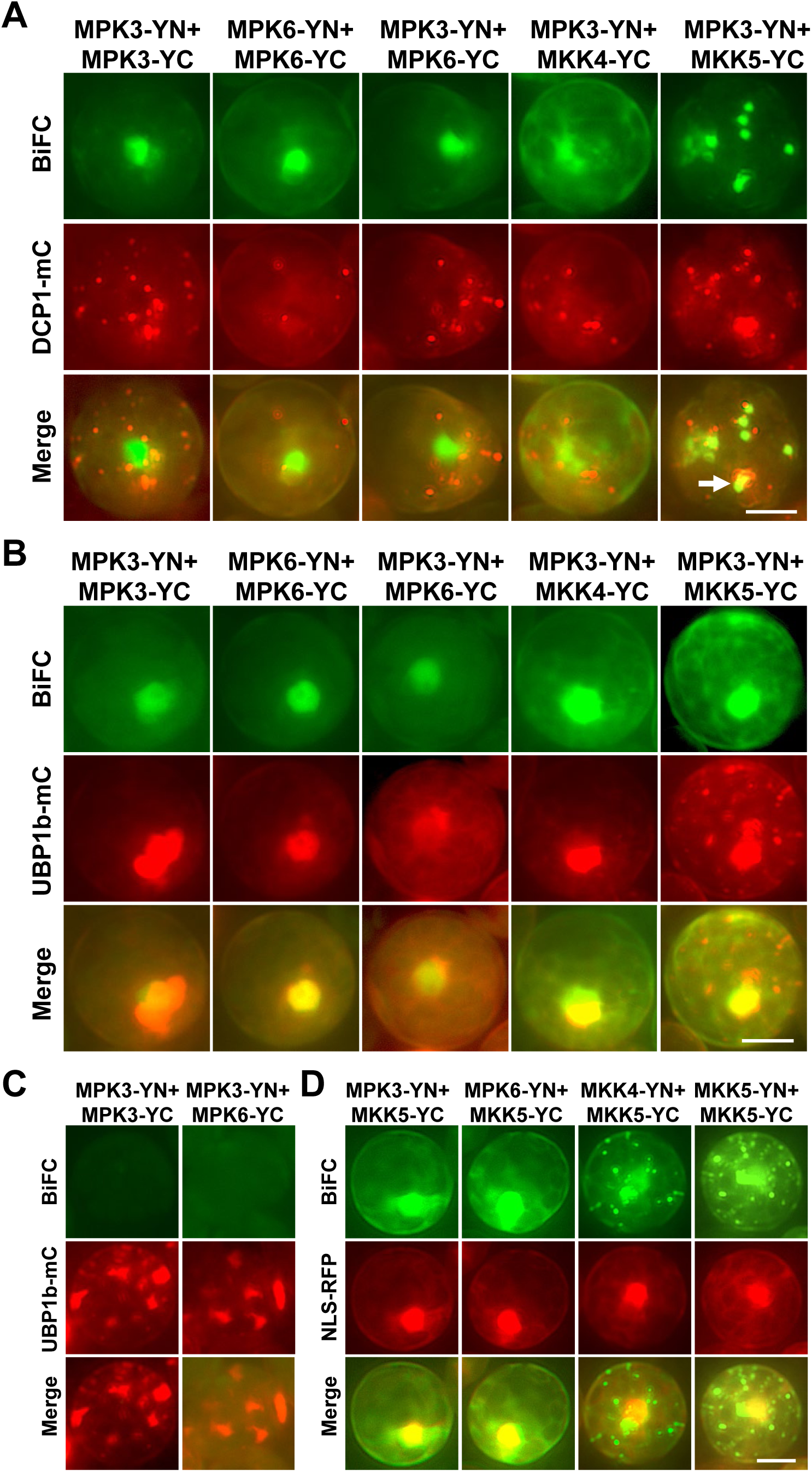
Protein-protein interactions of MPK3/6 and MKK4/5 are primarily taken place in SGs. **(A-B)** The signals from BiFC analysis were very sparsely co-localized with PB marker DCP1- mCherry (merged panel, example indicated by an arrow), but completely co-localized with SG marker UBP1b-mCherry (merged panel). **(C)** BiFC signals involved MPK3 were significantly diminished when co-expressed with UBP1b-mCherry. (**D)** The BiFC signals of MPK-MKK were primarily localized in the nucleus, whereas MKK-MKK signals were in cytoplasmic foci. Scale bar= 10 μm.

To determine if MPK3/6 and MKK4/5 self- and cross-interactions were taken place in PBs or SGs, the respective marker was co-expressed in the BiFC analyses. Results indicated that most of these interactions were not colocalized with a major PB component DCP1 (Fig. 2A), whereas almost completely co-localized with SG marker UBP1b (Fig. 2B). Note that BiFC signals of MPK3-MKK5 were occasionally overlapped with DCP1-mCherry signals (indicated by an arrow in Fig. 2A). Furthermore, UBP1b could be localized to both SGs and the nucleus hence some of these interactions such as MPK3 self-interaction and MPK3-MPK6 cross-interaction were co- localized with UBP1b in the nucleus (Fig. 2B). Intriguingly, the above-mentioned interactions appeared to be suppressed by the co-expression of UBP1b in some cells, as evidenced by very weak or missing BiFC signals and strong UBP1b-mcherry signals in SGs (Fig. 2C). This raised a possibility that MPK3-MPK3 and MPK3-MPK6 interactions were more stable in the nucleus than in SGs (with co-expressed UBP1b). For MKK4 and MKK5, the BiFC signals of both self- and cross-interactions were predominately localized in cytoplasmic granules and only partially (indicated by arrows) co-localized with DCP1, but almost completely co-localized with SG marker UBP1b (Fig. 3).

**Figure 3.**
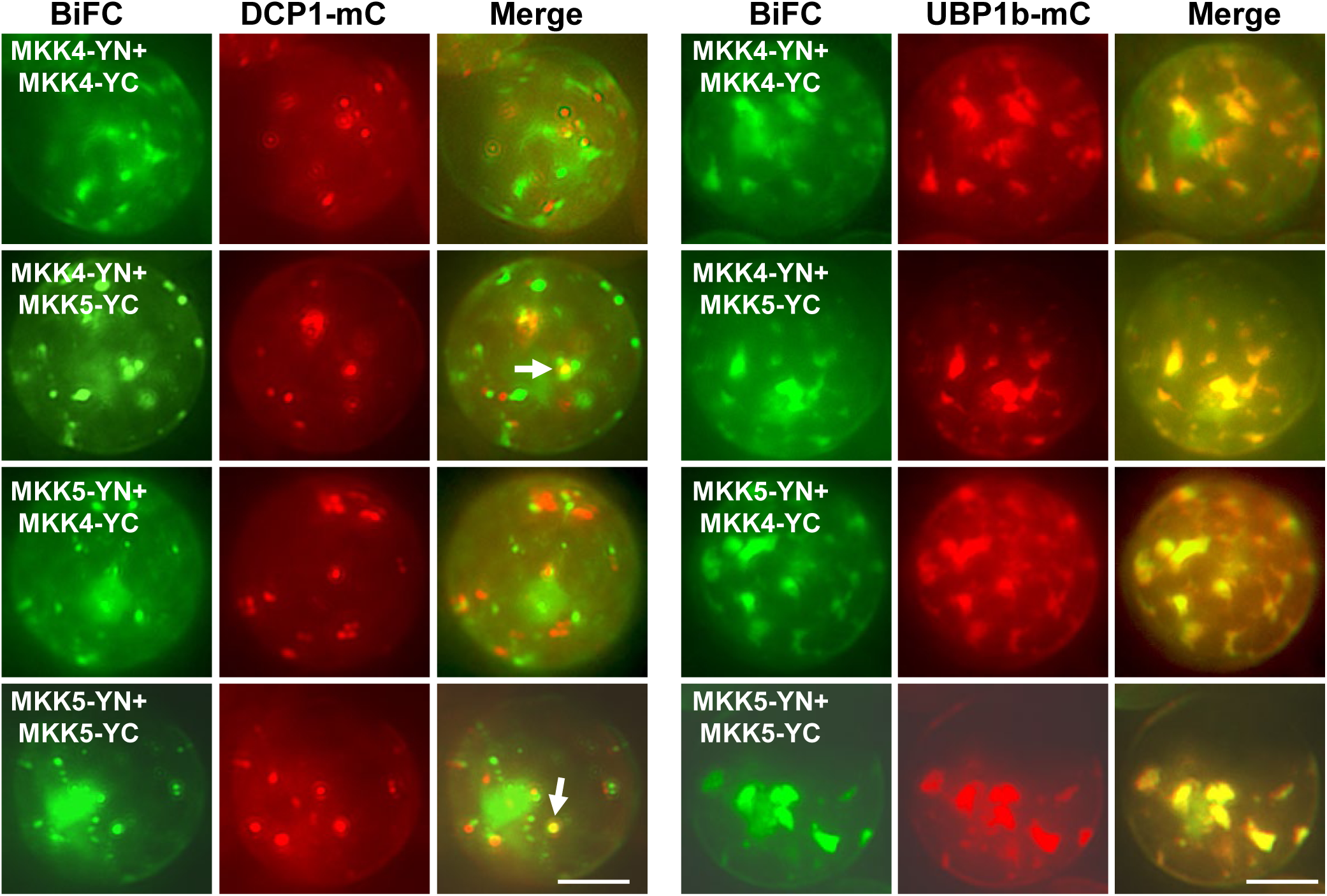
**MKK4 and MKK5 self- and cross-interactions are taken place primarily in SGs in BiFC analyses.** The BiFC signals were very sparsely co-localized with PB marker DCP1-mCherry (left panel, examples indicated by arrows), but completely co-localized with SG marker UBP1b-mCherry (right panel). Scale bar= 10 μm.

To validate protein-protein interactions identified in BiFC analyses, co-immunoprecipitation (Co-IP) assays were conducted. Pair-wise protein components with various tags were co-expressed in an Arabidopsis protoplasts transient expression system and immunoprecipitation was carried out using GFP-antibody. Results indicated that MKK4 and MKK5 (Fig. 1C) as well as MPK3 and MPK6 (Fig. 1D) could self- and cross-interact. MKK4 or MKK5 could cross-interact with MPK3 or MPK6 as well (Fig. 1E). No Co-IP signals were found when each of the MKK or MPK constructs was co-expressed with the empty construct with free GFP (Fig. 1F), indicating the specificity of the Co-IP results. Based on the results of BiFC (Fig. 1A-B and 2-3), Co-IP (Fig. 1C-F), and previous report (He *et al*., 2024), we proposed a model in which TZF1, MPK3/6, and MKK4/5 formed an interactome in SGs (Fig. EV1).

### DCP1 granule assembly affected by MPK3/6 and MKK4/5

In the course of conducting BiFC analyses, it was noted that the signal levels of PB component DCP1-mCherry varied, depending on specific co-expressed pair of BiFC constructs. For example, the DCP1 granule number was abundant when co-expressed with MKK4- nYFP+MKK4-cYFP, but much lower when co-expressed with MKK4-nYFP+MKK5-cYFP, MKK5-nYFP+MKK4-cYFP, and MKK5-nYFP+MKK5-cYFP, respectively (Fig. EV2). Likewise, DCP1 granule number was abundant when co-expressed with MPK3-nYFP+MPK3- cYFP, but low when co expressed with MPK3-nYFP+MKK4-cYFP, MPK3-nYFP+MKK5-cYFP, and MPK3-nYFP+MPK6-cYFP, respectively (Fig. EV3). It was also noted that both the number and the size of DCP1 granules were varied among samples. To verify if differential signal levels of DCP1 granules were due to individual proteins, additional co-expression analyses were conducted. The number of DCP1 granules was moderately suppressed by co-expression of MPK6 but severely suppressed by MKK5, as compared to MPK3 and MKK4, respectively. The size of DCP1 granules appeared to be suppressed by MKK4/5, but not MPK3/6 (Fig. 4A-B). Given MPK6 appeared to have a stronger effect than MPK3, additional family members of MPKs were tested using available BiFC constructs. Results showed that the BiFC constructs were as effective as the GFP-fusion constructs, because MPK6-cYFP could suppress DCP1 granule accumulation, as compared to MPK6-GFP. In additional to MPK6, MPK1, 4, and 11 had similar effects in suppressing DCP1 granule accumulation (Fig. EV4).

**Figure 4.**
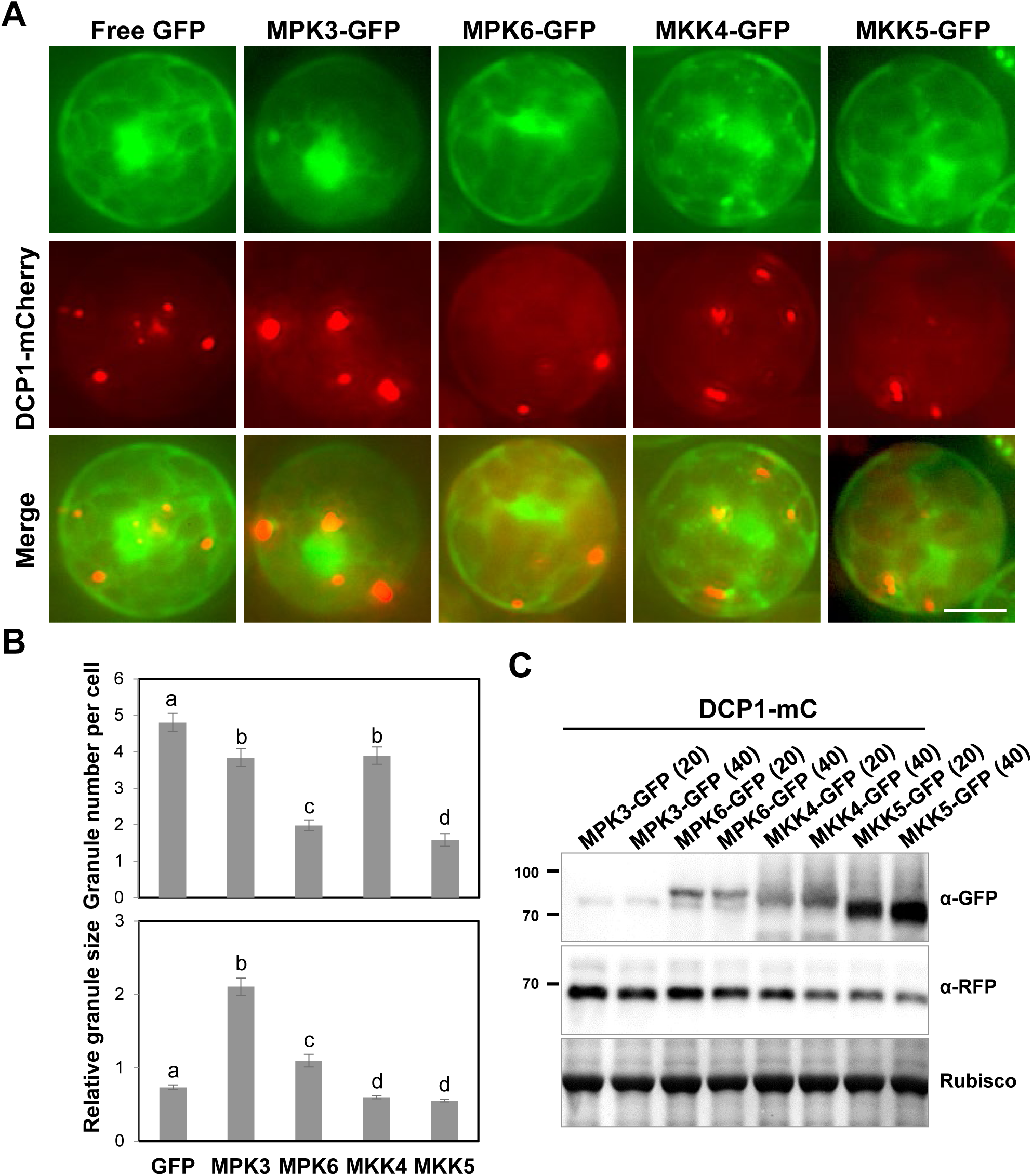
MPK3/6 and MKK4/5 affect the number and size of DCP1-mCherry granules. **(A)** Paired plasmid DNA constructs as indicated were co-expressed in Arabidopsis protoplasts transient expression analysis. Shown are free GFP, MPK3/6-GFP, MKK4/5-GFP (green signals), DCP1-mCherry (red signals), and merged images. All images were taken with the same exposure times. Scale bar= 10 μm. **(B)** Quantitative analysis of granule number per cell (upper panel) and average granule size (lower panel) as shown in (A). Columns represent means ± *SE*. Different letters above the bars indicate significant differences determined by ANOVA (*P* < 0.05). **(C)** DCP1-mCherry was co-expressed with two different doses (20 vs 40 mg of plasmid) of MPK3/6-GFP or MKK4/5-GFP in a protoplast transient expression assay. Immunoblot analysis was conducted using protein samples as indicated.

From the cellular images in Fig. EV4, it appeared that the protein expression levels of MPKs, MKKs and DCP1 varied in different samples. As protein level might affect biomolecular condensate assembly (Liu *et al*, 2023), immunoblot analyses were conducted. To further delineate the effects of MPKs/MKKs on DCP1 accumulation, two dozes of plasmid DNA (20 and 40 μg) were used in the protoplast transient expression analysis. Consistent with cellular images in Fig. EV4, the levels of protein expression displayed large differences ranging from weak to strong in the order of MPK3, MPK6, MKK4, and MKK5. By contrast, DCP1 level varied just slightly between samples co-expressed with MPK3/6 and MKK4/5, with only negligible differences between the two samples transformed with different doses of plasmid DNA (Fig. 4C). Similar analysis was carried out for various MPK-cYFP and DCP1-mCherry samples as shown in Fig. EV5. Results showed whereas the levels of MPKs varied, with MPK11-cYFP at the lowest abundance, the level of DCP1-mCherry remained at similar levels across different samples. Of note, MPK3-cYFP and MPK6-cYFP appeared to be much more stable than their counterparts fused with GFP. However, the co-expressed DCP1-mCherry was accumulated at a remarkably consistent level across all samples. Together, these results suggest that post-translational regulation plays a major role on DCP1 granule dynamics, although DCP1 protein accumulation might also play a secondary role.

### Phosphorylation activity of MKK5 modulates the partition of DCP1 between PBs and SGs

We have shown previously that plant SG assembly is affected by the phosphorylation status of the core proteins (He *et al*., 2024). Co-expression analysis was then conducted to determine if the phosphorylation status of MKK5 could affect these processes. Results showed that co-expression of MKK5^WT^ reduced the number of DCP1 granules to about 50%, whereas the MKK5^DD^ (a constitutively active form of MKK5^T215D/S221D^ (Ren *et al*, 2002; Zhao *et al*, 2017) exerted even stronger suppression, in contrast to the MKK5^KR^ (loss-of function mutation of the conserved K to R change in the kinase ATP-binding loop of MKK5^K99R^) (Ren *et al*., 2002) that enhanced the accumulation of DCP1 granules (Fig. 5A-B). Noticeably, the decrease of DCP1 granules was associated with an increased number of cells containing a dominant large DCP1 granule. We have shown previously that flg22-induced MAPK signaling cascade could trigger DCP1 phosphorylation and rapid disassembly of DCP1 granules, whereas the phospho-dead DCP1^S237A^ was resistant to the effect of flg22 (Yu *et al*, 2019). Here, by expressing DCP1 phosphorylation mutant constructs alone, it was found that the accumulation of granules from DCP1^S237A^ was far greater, whereas from DCP1^S237D^ (phospho-mimetic) was far less than that from DCP1^WT^ (Fig. 5). When the individual DCP1 constructs were co-expressed with phosphorylation mutants of MKK5, the DCP1^S237A^ granules remained highly abundant across each combination (Fig. 5C-D), whereas the accumulation of DCP1^S237D^ granules was low and dominated by the single large granule pattern across each combination of DCP1+MKK5 (Fig. 5E-F). These results suggest that MKK5 acts upstream of DCP1 and phosphorylation status of DCP1 is a determinant for the accumulation/assembly and form/size of DCP1 granules.

**Figure 5.**
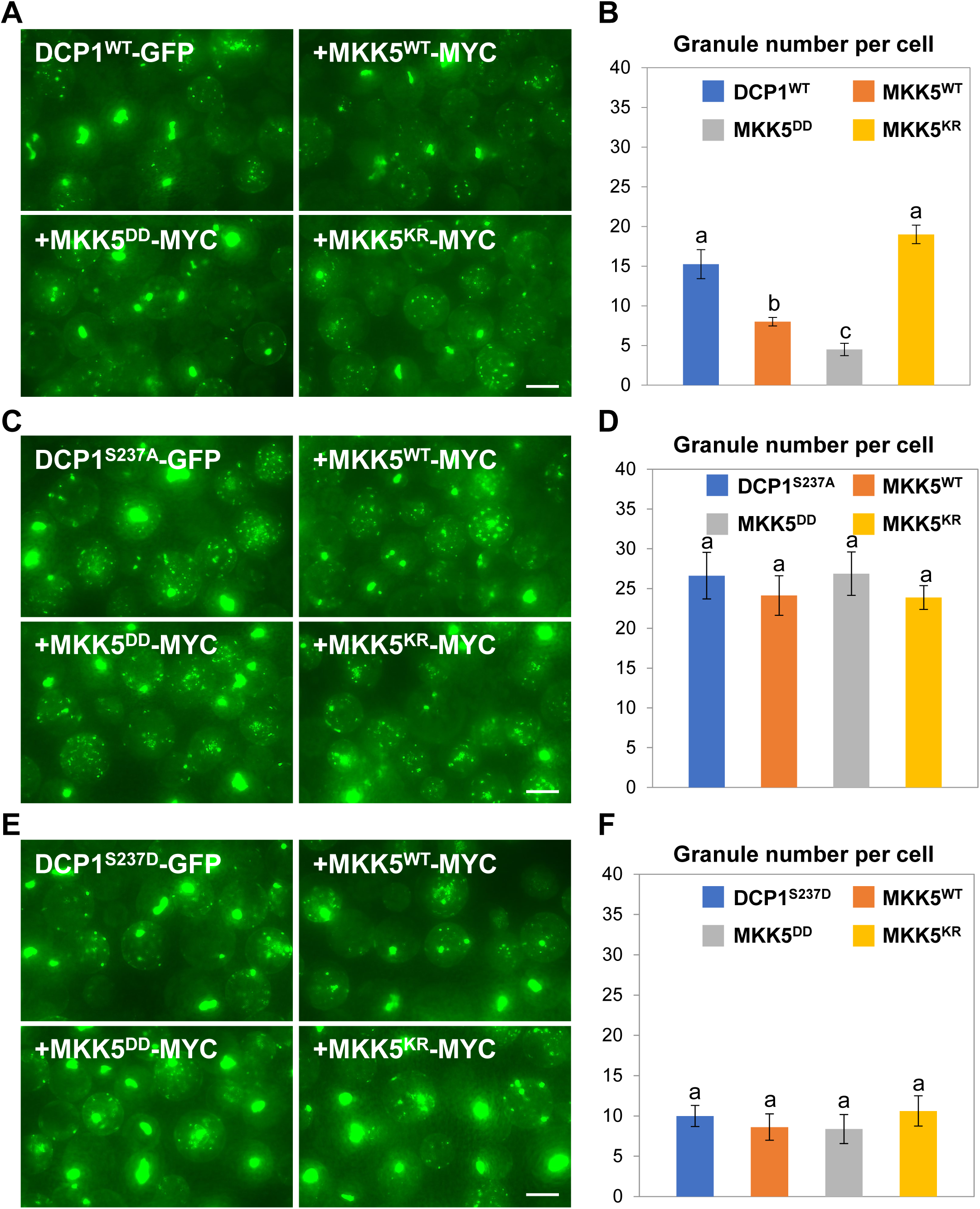
MKK5 kinase activity affects DCP1-GFP granule dynamics. **(A)** The number of typical (small) DCP1^WT^ -GFP granules was reduced by co-expression of the constitutive active MKK5^DD^, but increased by the constitutive inactive MKK5^KR^. Conversely, the number of atypical (large) DCP1^WT^-GFP granules was increased by co-expression of the constitutive active MKK5^DD^, but reduced by the constitutive inactive MKK5^KR^. Scale bar= 20 μm. **(B)** Quantitative analysis of typical small granule number per cell as shown in (A). Columns represent means ± *SE*. Different letters above the bars indicate significant differences as indicated by ANOVA (*P* < 0.05). **(C)** The number of typical (small) DCP1^S237A^-GFP granules was greater than that of the DCP1^WT^-GFP, and was relatively unaffected by co-expression of the MKK5^WT^, constitutive active MKK5^DD^, or the constitutive inactive MKK5^KR^. Scale bar= 20 μm. **(D)** Quantitative analysis of typical small granule number per cell as shown in (C). Columns represent means ± *SE*. Different letters above the bars indicate significant differences as indicated by ANOVA (*P* < 0.05). **(E)** The number of typical (small) DCP1^S237D^-GFP granules was smaller than that of the DCP1^WT^-GFP, and was relatively unaffected by co-expression of the MKK5^WT^, constitutive active MKK5^DD^, or the constitutive inactive MKK5^KR^. Scale bar= 20 μm. **(F)** Quantitative analysis of typical small granule number per cell as shown in (E). Columns represent means ± *SE*. Different letters above the bars indicate significant differences as indicated by ANOVA (*P* < 0.05).

To determine if MKK5-mediated DCP1 granule dynamics was related to protein accumulation, immunoblot analysis was conducted. Interestingly, compared to the MKK5^WT^, MKK5^DD^ was accumulated at a higher level, whereas MKK5^KR^ was accumulated at a lower level, across various samples. By contrast, the accumulation of DCP1^WT^, DCP1^S237A^, and DCP1^S237D^ showed a remarkably consistent level across various samples (Fig. 6). These results indicate that DCP1 granule dynamics is likely orchestrated by post-translational regulation but not protein accumulation. As DCP2 granule dynamics was similarly regulated by MKK5 kinase activity, same assay was also conducted. Similar results were obtained as those from DCP1 (Figure 6, right panel).

**Figure 6.**
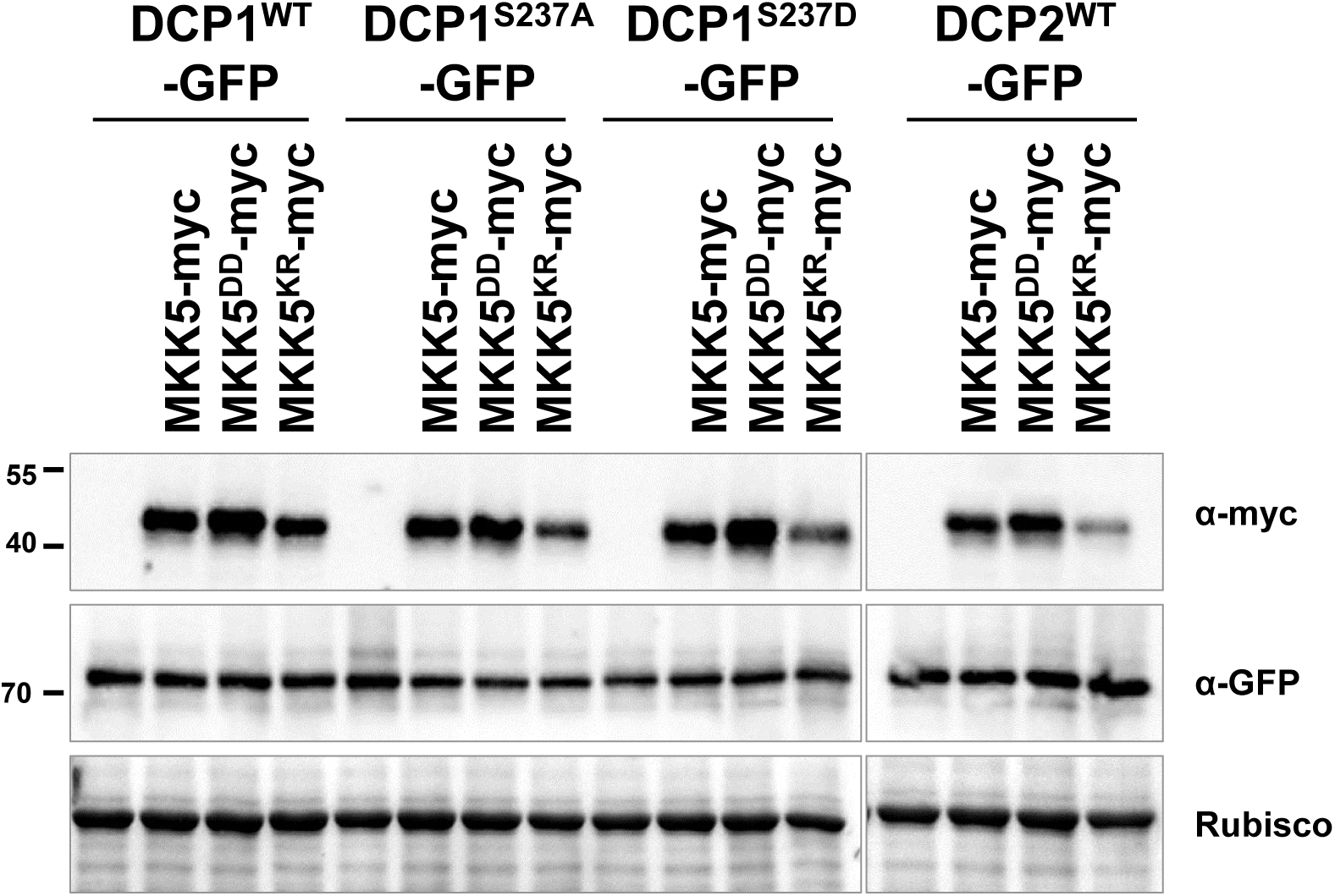
**The accumulation of DCP1-GFP and DCP2-GFP is relatively unaffected by the kinase activity of MKK5.** DCP1^WT^-GFP, DCP1^S237A^-GFP, DCP1^S237D^-GFP, or DCP2^WT^-GFP was co-expressed with various types of MKK5^WT, DD, KR^ in a protoplast transient expression assay as shown in Figure 5. Immunoblot analysis was conducted using protein samples as indicated.

### Phosphorylation triggers the sequestration of DCP1 from PBs to SGs

As a striking high number of cells contained seemingly coalesced large DCP1^S237D^ granules in Fig. 5E, we sought to determine the identity of the large DCP1 granules. As shown in Fig. 5, albeit differing in number, both large and small granules were present in all three types of DCP1, independent of its phosphorylation status. We first examined the relationship between small granules and various cellular markers by selecting the cells with desirable patterns. Results showed that small granules of all 3 types of DCP1 were independent of nuclear marker NLS-RFP. The small granules of DCP1^WT^ and DCP1^S237A^ were not co-localized with the SG marker UBP1b, supporting their identity as PBs. By contrast, the DCP1^S237D^ granules were generally larger and they completely co-localized with UBP1b, suggesting that phosphorylation of DCP1 is a trigger to sequester DCP1 from PBs to SGs (Fig. 7A).

**Figure 7.**
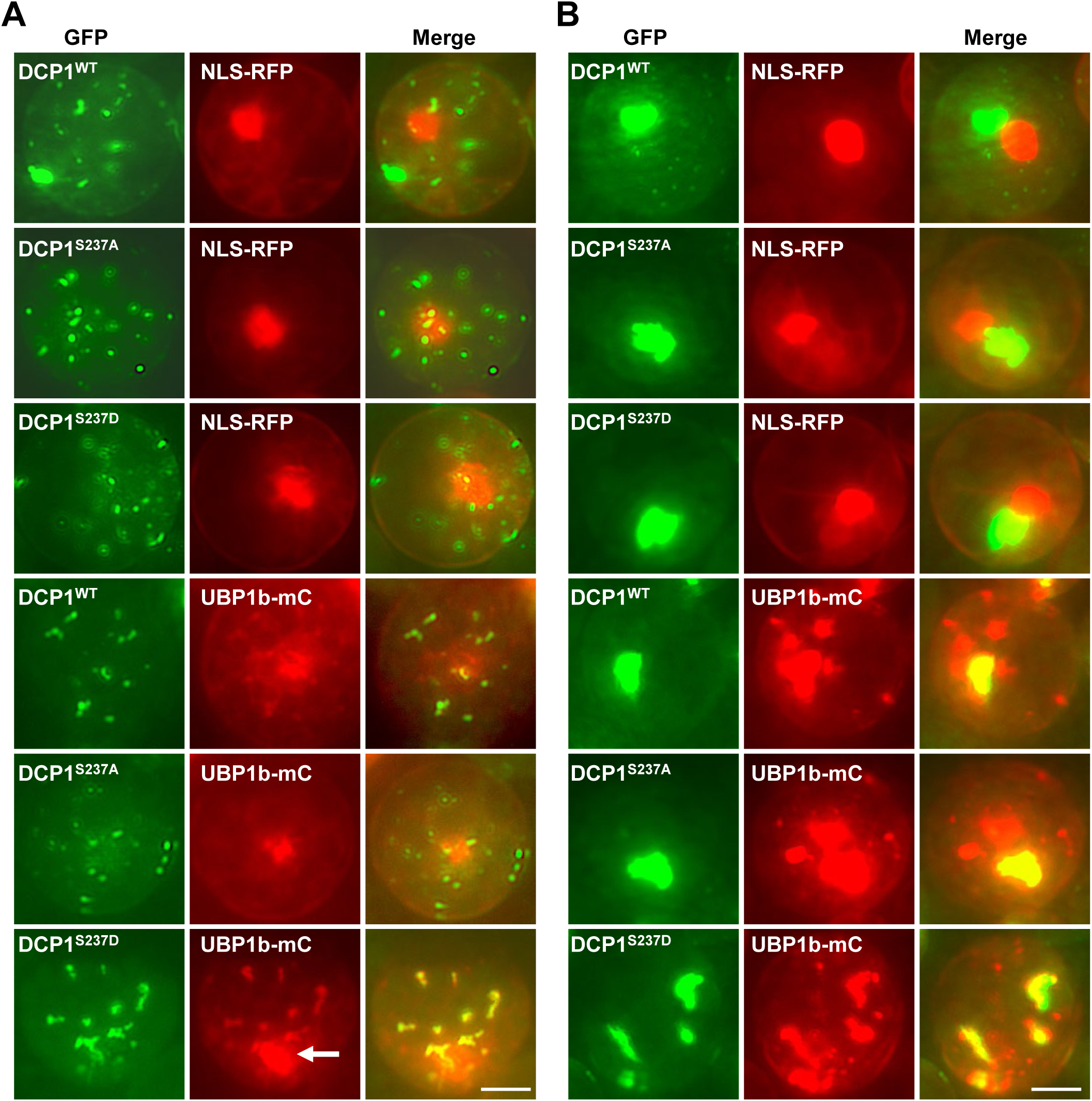
Co-localization of DCP1 granules with SG marker UBP1b. **(A)** In the cells with typical small PB-like DCP1 granules, whereas DCP1^WT^ and DCP1^S237A^ granules are largely independent of, the larger DCP1^S237D^ granules are colocalized with SG marker UBP1b-mCherry. The largest UBP1b-mC granule is the nucleus (arrow). **(B)** The coalesced large DCP1 granules are not co-localized with the nuclear marker NLS-RFP but are partially or completely co-localized with the SG marker UBP1b-mCherry. In this scenario, phosphorylation status of DCP1 does not affect the sub-cellular localization, because large granules from DCP1^WT^, phospho-dead (DCP1^S237A^), or phospho-mimetic (DCP1^S237D^) can still co-localize with UBP1b-mCherry. Scale bar= 10 μm.

Paradoxically, the large ‘nucleus-like’ DCP1^WT^ granules were not co-localized with the nuclear marker NLS-RFP, but instead partially or completely co-localized with the SG marker UBP1b. Similar scenarios were found for the large DCP1^S237A^ or DCP1^S237D^ granules (Fig. 7B). These results suggest that phosphorylated DCP1 is mainly sequestered to SGs, but the phosphorylation is not the only prerequisite for SG localization, as fewer cells with large granules were also found in DCP1^S237A^ samples and they could also co-localize with UBP1b. We speculate that there might be redundant post-translational modification events that could trigger the sequestration of DCP1^S237A^ to SGs. Together, these results suggest that protein phosphorylation and additional post-translational modification mechanisms likely play a role in the sequestration of DCP1 between PBs and SGs.

### UBP1b suppresses MAPK signaling to maintain DCP1 in PBs

In contrast to the relationship between DCP1 and MPK3/6 and MKK4/5, the co-expressed UBP1b appeared to affect the BiFC signals generated from various interactions within and between MPK3/6 and MKK4/5. For example, the BiFC signals from MKK4-nYFP+MKK4- cYFP and MKK5-nYFP+MKK4-cYFP were dampened by the co-expression of UBP1b-mCherry (Fig. EV6). In addition, the BiFC signals from MPK3-nYFP+MPK3-cYFP and MPK3- nYFP+MPK6-cYFP were also reduced by the co-expression of UBP1b-mCherry (Fig. EV7). These results were intriguing because while it seemed to be clear that BiFC signals of MKK5 self-interaction was unaffected, the BiFC signals of MKK5-nYFP+MKK4-cYFP were suppressed (Fig. EV6). Likewise, it was unclear why the BiFC signals of MPK3 self-interaction and MPK3-nYFP+MPK6-cYFP were suppressed, but not for MPK3 interaction with either MKK4 or MKK5 (Fig. EV7). To clarify if UBP1b suppressed the signals of protein-protein interactions or the accumulation of individual protein itself, co-expression analyses were conducted using single protein constructs. Results showed that the signals from MPK3 and MKK4 were strongly, and from MPK6 was modestly dampened by the co-expression of UBP1b (Fig. 8A). Consistent with the cellular imaging results, immunoblot analysis revealed that co-expression of two different doses of UBP1b resulted in a reduction of the accumulation of MPK6 and MKK4, with a lesser extent on MKK5. MPK3 level was too low to be determined (Fig. 8B). Given UBP1b appeared to cause a general reduction of MPKs/MKKs protein accumulation, the variation of MPKs/MKKs granule intensity/dynamics could have been contributed by the protein abundance, although it was not as obvious for MPK6 and MKK5 from cellular images (Fig. 8A).

**Figure 8.**
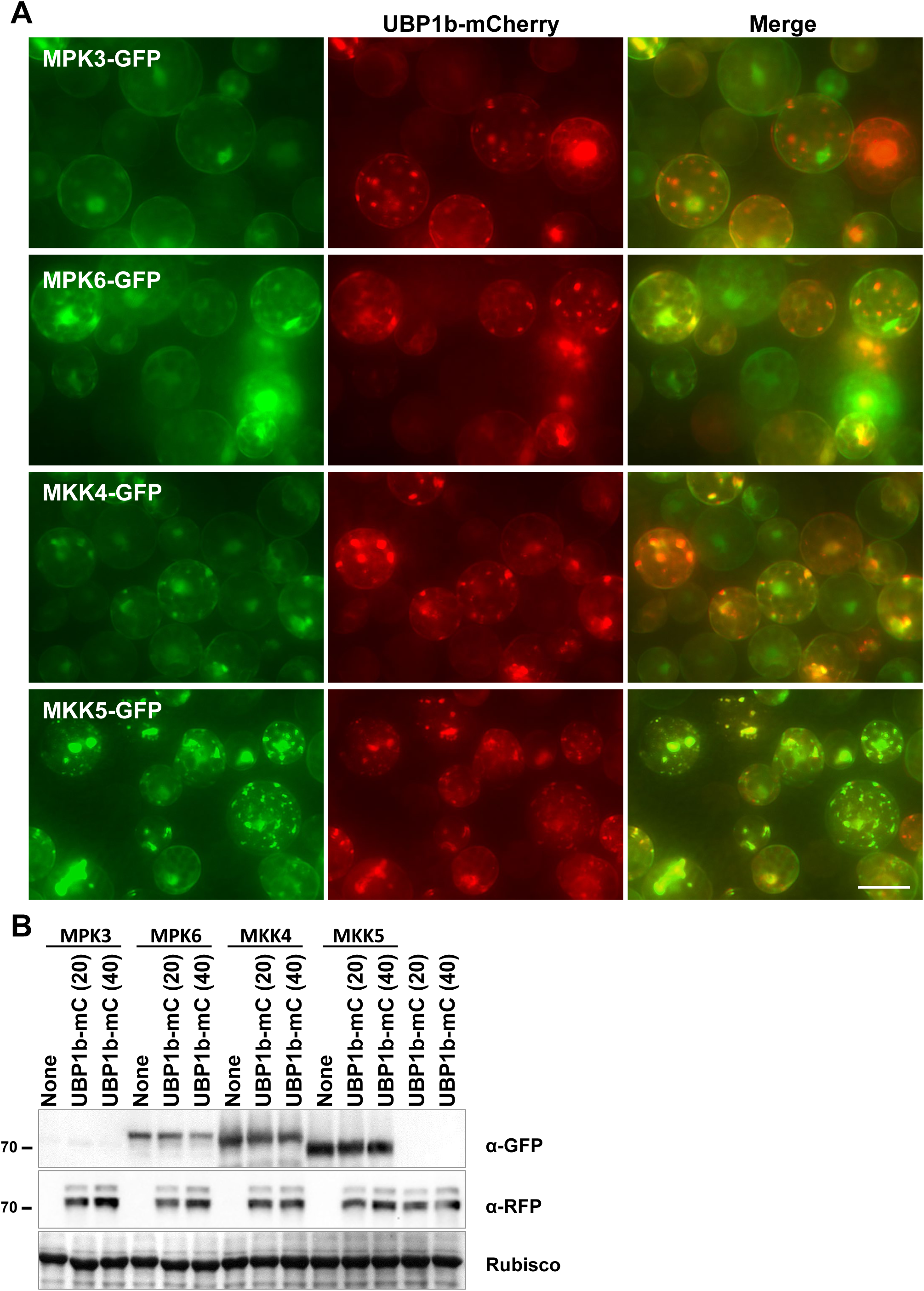
The SG marker UBP1b-mCherry suppresses the expression of MPK3 and MKK4 when co-expressing in an Arabidopsis protoplast transient expression analysis. **(A)** The expression of MPK3-GFP and MKK4-GFP appeared to be suppressed. All the green and red images were taken with the same exposure time. Scale bar= 20 μm. **(B)** Immunoblot analysis to determine protein accumulation from experiment similar to what is shown in (A).

Given DCP1 granule assembly was affected by MAPK signaling components whose accumulation appeared to be controlled by UBP1b, the relationship between DCP1 phosphorylation status and UBP1b was investigated. Compared to the DCP1^WT^, DCP1^S237A^ had increased and DCP1^S237D^ had decreased number of granules (Fig. 9A). When UBP1b was co-expressed, DCP1 granule abundance was enhanced and there was no significant difference between the three types of DCP1 (Fig. 9B-C). Because none of the DCP1-mCherry granules were co-localized with the SG marker UBP1b-GFP, the small and distinct DCP1-mCherry granules were likely PBs (Fig. 9B). It is worth mentioning that DCP1^S237D^-GFP granules were completely co-localized with the SG marker UBP1b-mCherry (Fig. 7A). This was due to the use of minimal amount of UBP1b-mCherry plasmid in that experiment and UBP1b-mCherry was accumulated at a much lower level than that of UBP1b-GFP used in this experiment. As UBP1b- mCherry accumulation did not reach the threshold level to trigger DCP1 sequestration to PBs, the DCP1^S237D^-GFP granules were still maintained in SGs (Fig. 7A). Immunoblot analysis was conducted to determine if increased DCP1 granule abundance was correlated with higher level of protein accumulation. Compared to the samples co-expressed with free GFP, a general reduction of the accumulation of all three types of DCP1, independent of their phosphorylation status, was found when co-expressed with UBP1b-GFP. More importantly, the three DCP1 proteins accumulated at nearly the same level, ruling out the possibility of increased DCP1 granule abundance was due to elevated protein accumulation (Fig. 9D). Together, these results suggest that UBP1b suppresses MAPK signaling components that act as negative regulators of DCP1 granule assembly. We propose a model in which flg22 activates MAPK signaling mediated post-translational modification of DCP1 that results in a decrease of DCP1 localization to PBs, whereas an increase of DCP1 sequestration to SGs. UBP1b, on the other hand, acts as a negative regulator of certain MPKs and MKKs, hence counteracting with the MAPK signaling effects and resulting in the maintenance of DCP1 to be associated with PBs. Because UBP1b-mediated increase of DCP1 sequestration to PBs appeared to override the phosphorylation status of DCP1, it is likely that an additional unknown mechanism exerted by UBP1b is also operating in this process. We also speculate that UBP1b, a key modulator of SG assembly, and MAPK signaling play a role in the homeostasis control of PBs and SGs in response to stresses other than flg22 (Fig. 10), such as heat, salt, and ABA (Nguyen *et al*, 2016; Nguyen *et al*, 2017; Yan *et al*, 2022).

**Figure 9.**
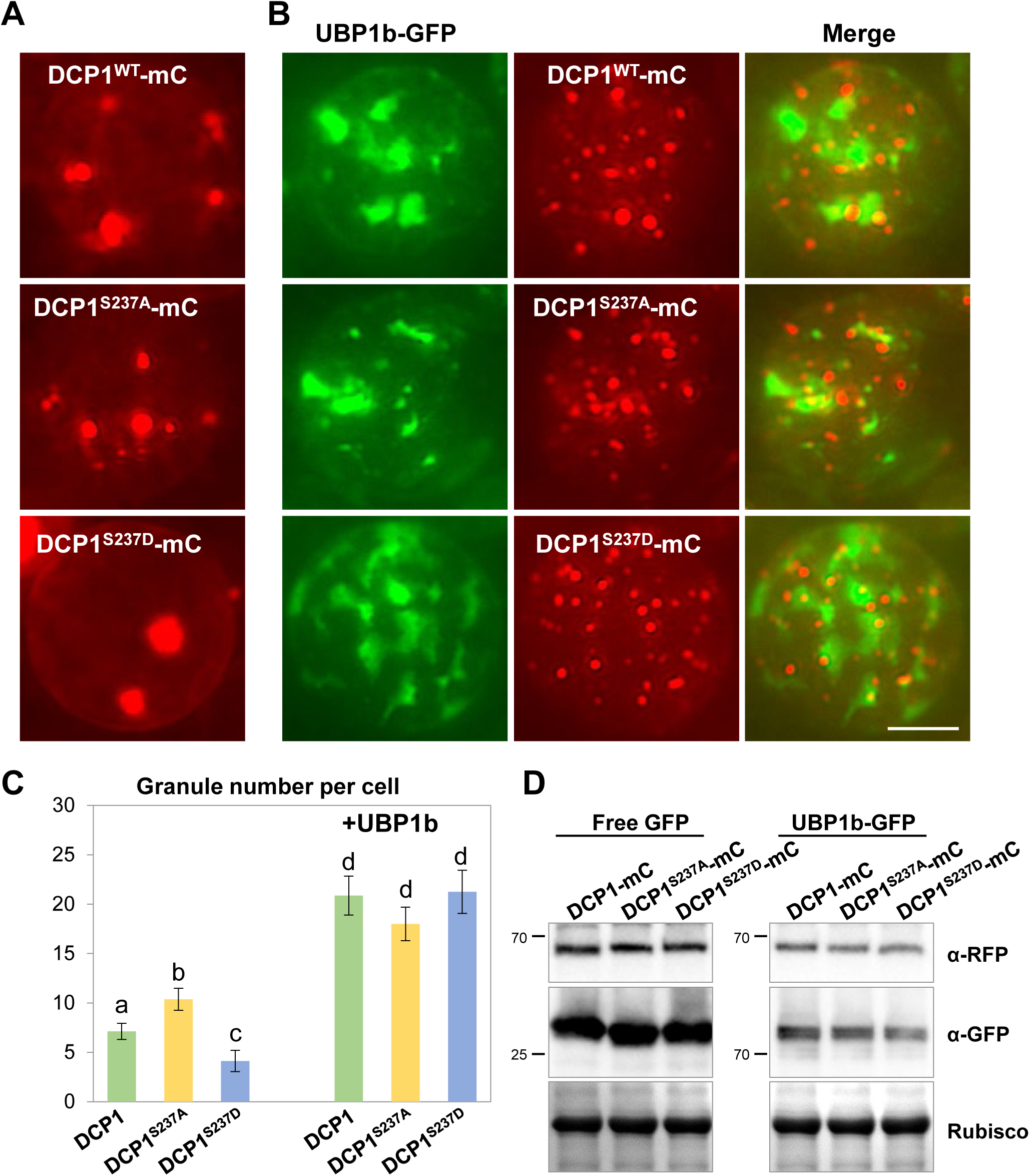
The SG marker UBP1b-GFP enhances DCP1-mCherry granule assembly when co-expressed in an Arabidopsis protoplast transient expression analysis. **(A)** DCP1-mCherry granule assembly in the absence of UBP1b-GFP. **(B)** UBP1b-induced enhancement of granule assembly is independent of DCP1 phosphorylation status as neither phospho-dead DCP1^S237A^ nor phospho-mimetic DCP1^S237D^ is different from DCP1^WT^. All the images were taken with the same exposure time. Scale bar= 10 μm. **(C)** Quantitative analysis of typical small granule number per cell as shown in (A) and (B). Columns represent means ± *SE*. Different letters above the bars indicate significant differences as indicated by ANOVA (*P* < 0.05). **(D)** Immunoblot analysis to determine DCP1-mChery accumulation in the presence of free GFP vs UBP1b-GFP.

**Figure 10.**
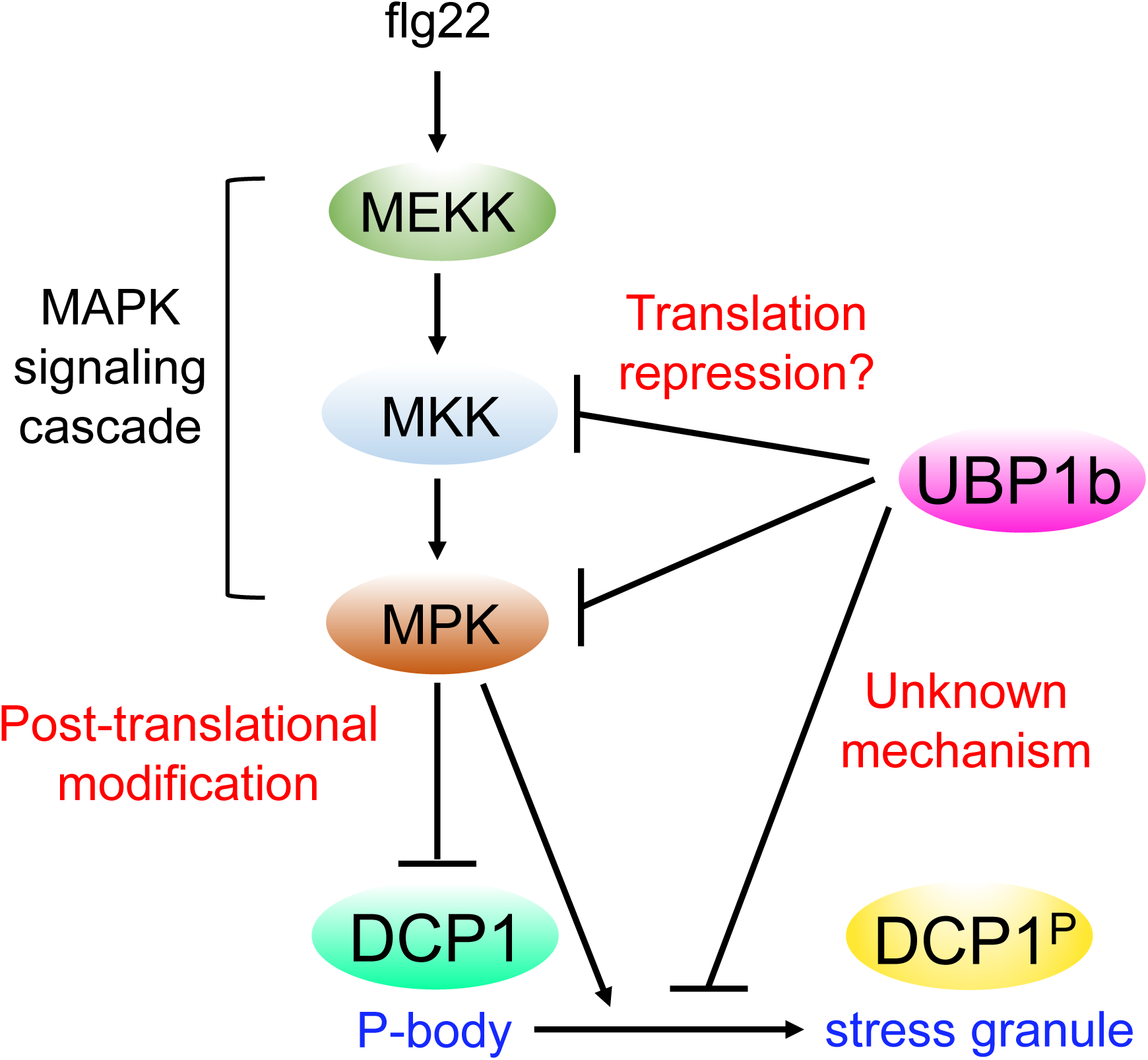
**Working model of current study.** The bacterial flg22, a pathogen-associated molecular pattern, triggers the innate immune response via MAPK signaling cascade. By an unknown mechanism, this activation causes a quick and transient disappearance of DCP1 granules. In this study, we show that the MPK3/6 and MKK4/5 form a protein interacting network (Figure EV1) in SGs and the kinase activity of MAPK cascade is required to suppress DCP1 localization to PBs, while promote DCP1 to be associated with SGs. On the other hand, the SG marker UBP1b can suppress the accumulation of MPK3/6 and MKK4/5 perhaps via mRNA binding and/or translation repression, hence diminishing the effect of MAPK signaling and maintaining DCP1 in PBs. UBP1b also mediates another unknown post-translational regulatory mechanism to maintain DCP1 in PBs.

## Discussion

PBs and SGs have intimate relationship in composition and function in various cellular processes. Although distinct core components of PB and SG have been redefined by using sophisticated proximity mapping (Youn *et al*, 2018), recent reports have also found an extensive overlap of composition across PBs and SGs, such as poorly translated mRNAs and low complexity RNA-binding proteins (Kershaw *et al*, 2021). In mammals, double knockout of RNA-binding protein G3BP1 and G3BP2 prevents SG assembly induced by eukaryotic initiation factor 2α phosphorylation. In this double mutant background, phosphor-mimetic mutant G3BP^S149E^ failed to rescue SG assembly, highlighting the importance of the role of PTM on G3BP. Caprin and USP10 bind G3BP in a mutually exclusive way, whereas G3BP-Caprin complex promotes SG assembly, and G3BP-USP10 complex promotes SG disassembly (Kedersha *et al*, 2016; Krapp *et al*, 2017). The plant ubiquitin-specific protease family members are homologs of USP10, including UBP24, a negative regulator of ABA signaling (Zhao *et al*, 2016). It is not known if UBP24 is involved in PB-SG interaction. Confusing in nomenclature, although UBP1b is not an UBP family member, it is a key modulator of SG assembly in plants (Yan *et al*., 2022). On the other hand, the mammalian DDX6 (Rck/p54), a major PB scaffold component, plays a key role in PB-SG interaction. DDX6 limits itself and other RNPs to be assembled into SGs. In the absence of DDX6, more RNPs are partitioned into SGs. Loss PB scaffold proteins such as DCP1 and DDX6 also causes reduction in PB growth and enhances incompletely assembled PBs docking with SGs to form hybrid granules with irregular shapes (Majerciak *et al*, 2023; Ripin *et al*, 2024). SG assembly provides a means for the temporal and spatial compartmentalization of signaling components critical for cell growth and defense response (Kedersha *et al*, 2013). In fission yeast, while it is not demonstrated that MAPK signaling components are sequestered to SGs, the high heat stress induced MAPK activation triggers the sequestration of upstream regulator PKC into SGs thereby deactivating MAPK hyperactivation induced cell death (Kanda *et al*, 2021; Sugiura, 2021). While some limited information is available from studies using non-plant models, the molecular mechanisms mediating PB-SG dynamics in plants are virtually unknown.

### MPK-MKK-TZF1 interactome in SGs

In this report, we have found that MAPK signaling components, including MPK3, MPK6, MKK4, and MKK5, can self- and cross-interact with each other (Fig. 1-3). The cellular sites of these interactions are mainly in SGs, with a slight chance in PBs due to limited co-localization with a core PB component DCP1. MPK3 and MPK6 self- and cross-interactions, as well as cross-interactions between MPK3/6 and MKK4/5 often take place in the nucleus where the SG marker UBP1b can also be localized when co-expressed with the MAPK BiFC constructs (Fig. 2). In contrast, MKK4 and MKK5 self- and cross-interactions are primarily taken place in cytoplasmic granules with complete co-localization with SG marker UBP1b and limited co- localization with the core PB component DCP1 (Fig. 3). More importantly, all interactions are co-localized with TZF1 cytoplasmic granules, including MPK3 and MPK6 self- and cross-interactions (Fig. 1A-B). As we have demonstrated in an earlier report that TZF1 interacts with MPK3, MPK6, MKK4, and MKK5 *in vivo* and *in vitro* (He et al., 2024), we propose that TZF1 forms an interacting network with MAPK components in SGs (Fig. EV1).

### MAPK signaling modulates DCP1 granule assembly

In the process of conducting BiFC analyses, it was noted that the proteins from BiFC constructs had significant interactions with co-expressed PB (DCP1) or SG (UBP1b) marker protein. For example, DCP1-mCherry granules were suppressed by all combinations except MKK4-nYFP+MKK4-cYFP (Fig. EV2) and MPK3-nYFP+MPK3-cYFP (Fig. EV3). Further analysis indicated that DCP1-mCherry granules were differentially affected by MAPK signaling components: (1) DCP1 granule size was reduced in the order of co-expression of MPK3, MPK6, MKK4, and MKK5; (2) DCP1 granule number was reduced by co-expression of MPK6 and MKK5, compared to their counterparts MPK3 and MKK4, respectively (Fig. 4). Additional analysis revealed that MPK1, MPK4, and MPK11 could also reduce the number of DCP1- mCherry granules (Fig. EV4). This is potentially important because the two MKK4/5-MPK3/6 and MEKK1-MKK2-MPK4 MAPK signaling cascades play opposing roles in plant cold response (Zhao *et al*., 2017). Our preliminary results indicated that MEKK1 could enhance DCP1 granule assembly (Fig. EV8A) in a DCP1 phosphorylation-independent manner (Fig. EV8B). Moving forward, it would be interesting to clarify how components in the MEKK1- MKK2-MPK4 cascade affects DCP1 granule assembly. In fact, it would be more important to compare how MKK4/5-MPK3/6 and MEKK1-MKK2-MPK4 cascades affect plant PB/SG assembly in general. In an earlier stage of the research, we speculated that DCP1 granule assembly might be affected by differential protein accumulation of co-expressed MAPK signaling components. A series of protein gel-blot analyses revealed that although the level of protein accumulation of MPKs and MKKs was in a wide range, the co-expressed DCP1-mCherry accumulated in a remarkable consistent level. These results suggest that the size and number of DCP1 granules are modulated by MAPK signaling post-translationally.

To potentiate the idea that MAPK signaling can modulate DCP1 granule assembly, the kinase activity of MKK5 was examined, as MKK5 exerted a strong effect on DCP1 granule assembly. Results showed that kinase activity was required for MKK5 to reduce the number of DCP1 granules (Fig. 5A-B). It was also found that MKK5 kinase activity could trigger the assembly of unusually large DCP1 granules, which was validated to be SGs by co-localization with UBP1b (Fig. 7). On the basis of these findings, we hypothesized that MAPK signaling induced DCP1 phosphorylation could reduce the sequestration of DCP1 into PBs but enhance it into SGs. To test this hypothesis directly, we used phospho-dead form of DCP1^S237A^ and phospho-mimetic form of DCP1^S237D^. The results indicated that neither DCP1^S237A^ nor DCP1^S237D^ were affected by the kinase activity of MKK5. Furthermore, compared to the DCP1^WT^, there was a significant increase of small DCP1^S237A^ granules (PBs) (Fig. 5C-D) and decrease of small but increase of large DCP1^S237D^ granules (SGs) (Fig. 5E-F). More importantly, the level of protein accumulation across DCP1^WT^, DCP1^S237A^, and DCP1^S237D^ co-expressed with MKKs of different phosphorylation capacity showed a remarkable consistent level (Fig. 8), again supporting the notion that the dynamics of DCP1 granule assembly is orchestrated by MAPK signaling mediated post-translational modification. On the other hand, we showed previously that flag22- induced MPK3/6 phosphorylation of DCP1 is required for the positive effects of DCP1-DCP2 complex on plant microbe associated molecular patterns (MAMPs)-triggered responses and immunity against pathogenic bacteria (Yu *et al*., 2019). Given the new findings present here, it would be informative to determine if phosphorylated DCP1 localization in SGs is a pre-requisite for plant immunity.

In a parallel example, we reported recently that Arabidopsis TZF1 recruits MPK3 and MPK6 to SGs and TZF1 is phosphorylated by MPK3/6. Interestingly, TZF1 is differentially phosphorylated and de-phosphorylated on various residues by flg22-activated MAPK signaling cascade. Mutations of different TZF1 phosphorylation sites could either enhance or reduce TZF1 granules. Remarkably, some of the phosphorylation mutations could similarly trigger the assembly of unusually large TZF1 granules (He et al., 2024). As TZF1 is mainly localized in SGs and only partially co-localizes with PB components such as DCP1 and DCP2 (Pomeranz *et al*, 2010). As in the case of DDX6 (Rck/p54) in mammals (Ripin *et al*., 2024), it would be interesting in the future to further delineate how reversible phosphorylation of different residues plays the same or opposing role in shuttling TZF1 between PBs and SGs or simply enhance or reduce the size of TZF1 SGs.

### UBP1b counteracts MAPK signaling to maintain DCP1 in PBs

In contrast to the negative effects of MAPK signaling on DCP1 sequestration into PBs, here we have found that the SG marker UBP1b appeared to play a positive role in maintaining DCP1 in PBs (Fig. 9). UBP1b is an RNA-binding protein and a plant SG marker. UBP1b is a homolog of mammalian TIA-1 and TIAR that can promote the sequestration of untranslated mRNAs into SGs (Kedersha *et al*, 1999). The UBP1b SGs are induced by heat in the UBP1b overexpression (OX) plants that are heat stress tolerant (Nguyen *et al*., 2016). It was proposed that UBP1b sequesters mRNAs encoding DnaJ heat shock protein and other stress-related proteins into SGs to achieve heat-tolerance. UBP1b OX plants are also ABA hypersensitive. Curiously, *MPK3*, *MKK4*, and *MKK9* were up-regulated, but their half-lives were unaltered in UBP1b OX plants, indicating that these mRNAs were not the direct targets of UBP1b (Nguyen *et al*., 2017). The subcellular localization of MAPK signaling components and their relationship with UBP1b remains unclear. In this report, we have observed that some combination of MPK3/6 and MKK4/5 expressed via BiFC constructs appeared to be suppressed by the co-expression of UBP1b (Fig. EV6-7). Further analysis validated that UBP1b could reduce protein accumulation of MPK3/6 and MKK4/5 (Fig. 9). Given MAPK signaling negatively regulate DCP1 sequestration into PBs, we hypothesized that UBP1b could revert this negative regulation by suppressing MAPK signaling. Our hypothesis was supported by the results in which DCP1 was highly sequestered into PBs independent of its phosphorylation status when UBP1b was co-expressed (Fig. 9). Interestingly, the enhancement of DCP1 PB assembly was not due to elevated DCP1 protein accumulation, suggesting that the dynamics of DCP1 shuttling between PBs and SGs is controlled by MAPK signaling mediated post-translational modifications (Fig. 10).

In summary, although the interaction between PBs and SGs and molecules exchange between the two have been well-documented in mammals (Riggs *et al*., 2020), the molecular details of these processes are unknown in plants. Our findings have revealed a molecular mechanism mediating PB-SG dynamics in plants. We have shown recently that TZF1 with two intrinsically disordered domains is able to recruit MAPK signaling components to SGs (He *et al*., 2024). We have found that TZF1-MPK3/6-MKK4/5 forms a protein-protein interacting network. DCP1, a core component of plant PBs, is phosphorylated by MPK3/6 and the phosphorylation triggers a rapid reduction of DCP1 partition into PBs (Yu *et al*., 2019). Here we have found that this reduction is concomitantly associated with the increase of DCP1 partition into SGs, hence establishing a role for MAPK signaling in mediating PB-SG dynamics in plants. Furthermore, we have found that plant SG marker protein UBP1b plays a role in maintaining DCP1 in PBs by suppressing the accumulation of MAPK signaling components. Together, we propose that MAPK signaling and UBP1b modulate the dynamics of PBs and SGs in plants.

## Materials and methods

### Protoplast transient expression analysis

Arabidopsis protoplasts transient expression analyses were conducted mainly as described (Yoo *et al*, 2007), with additional modifications as described (He *et al*., 2024).

### BiFC analysis

The CDS of MKK4, MKK5, MPK3, and MPK6 were cloned into pA7-YN (containing N-terminal half of YFP) and pA7-YC (containing C-terminal half of YFP) vector (Chen *et al*, 2006), respectively. Each pair of BiFC construct and an additional cellular localization marker were co-transformed into Arabidopsis protoplasts.

### Co-IP assay

Total proteins from Arabidopsis protoplasts co-expressing plasmid pairs were lysed with lysis buffer (100 mM Tris pH 8.0, 150 mM NaCl, 5 mM EDTA, 10 mM DTT, 0.1% NP-40, proteases inhibitor cocktail). Extracted proteins were then incubated with equilibrated GFP-trap beads (Chromotek) at 4°C for 2 hr under gentle agitation, followed by 3 times of washing with wash buffer (100 mM Tris pH 8.0, 150 mM NaCl, 5 mM EDTA, proteases inhibitor cocktail). Immunoblots were performed using α-GFP (Roche), α-FLAG antibodies (Sigma) or α-myc antibodies.

## Accession numbers

The accession numbers used are as follows: TZF1 (At2g25900), DCP1 (At1g08370), UBP1b (At1g17370), MEKK1 (At4g08500), MKK4 (At1g51660), MKK5 (At3g21220), MPK1 (At1g10210), MPK3 (At3g45640), MPK4 (At4g01370), MPK6 (At2g43790), MPK7 (At2g18170), and MPK11 (At1g01560).

## Acknowledgements

We would like to thank undergraduate researchers Hannah Furness, Gianni Giarrano, Alex Tarter, and Yu Wang for assistance in various experiments. This work was supported by the grants from National Science Foundation MCB-1906060 to JC. Jang, P. He, L. Shan, and Y. Wang; Ohio State University President’s Research Excellence Accelerator award, Ohio Agricultural Research and Development Center SEEDS Program #2018007, Ohio State University College of Food, Agricultural, and Environmental Sciences Internal Grant Program #2022014, and Ohio State University Center for Applied Plant Sciences Research Enhancement Grant to JC Jang.

## Author contributions

S.-L.H. and J.-C.J. conceived and designed the experiments; S.-L.H. performed most of the experiments; S.-L.H. and J.-C.J. wrote the manuscript; P.H., L.S., and Y.W. provided comments, tools, and reagents for the project.

---

**Figure EV1.**
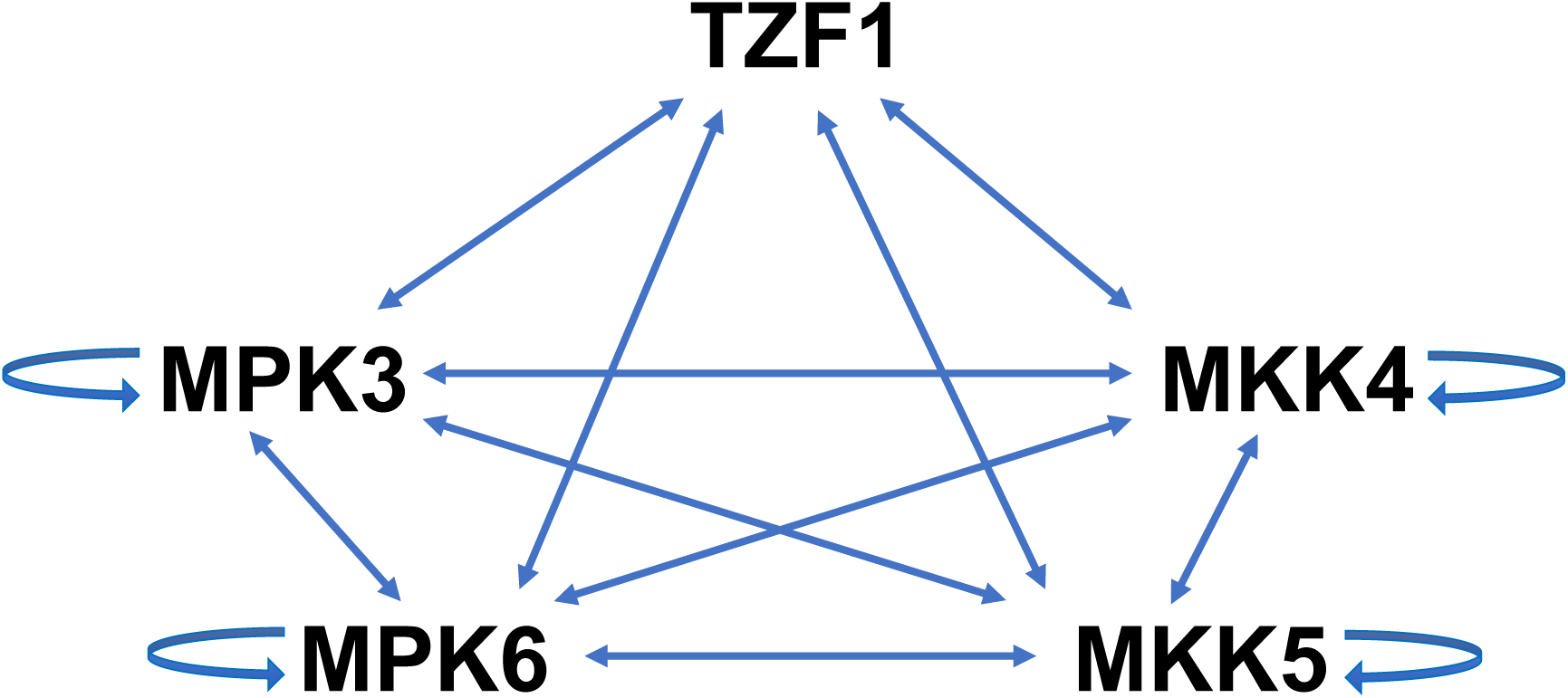
Proposed model of TZF1-MPK3/6-MKK4/5 interacting network in stress granules. The model is based on the results of current study and in a previous report (He *et al*., 2024) showing interaction between TZF1 and MPK3/6 and MKK4/5, respectively.

**Figure EV2.**
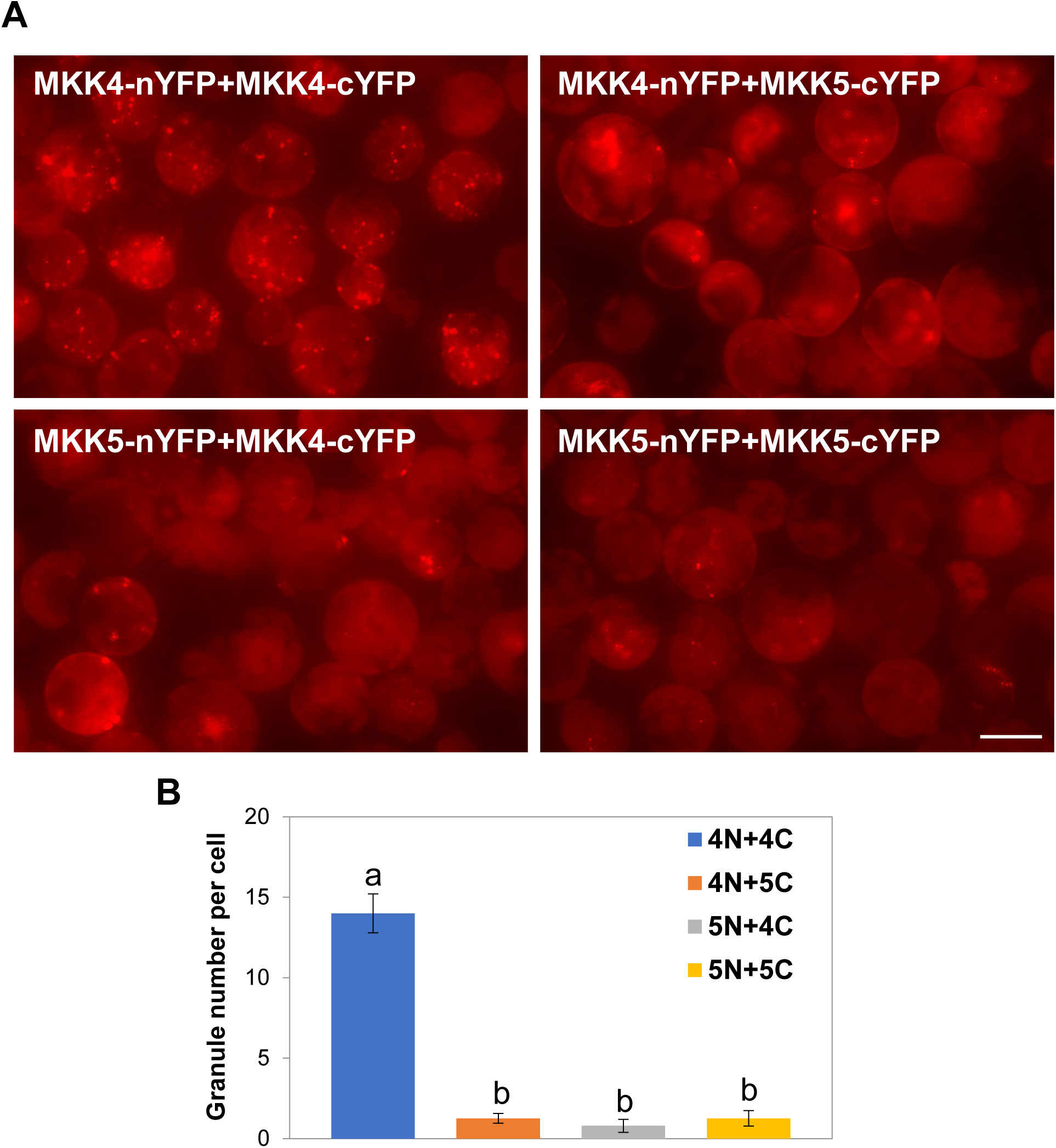
MKK4 and MKK5 interactions affect DCP1 granule assembly. **(A)** The granule assembly of co-expressed PB marker DCP1-mCherry is suppressed by hetero-dimers of MKK4 and MKK5 and homo-dimers of MKK5, whereas unaffected by homo-dimers of MKK4 (upper left), in BiFC analyses. Scale bar= 15 μm. Images of the BiFC signals (YFP) are not shown. **(B)** Quantitative analysis of granule number per cell as shown in (A). Columns represent means ± *SE*. Different letters above the bars indicate significant differences as indicated by ANOVA (*P* < 0.05).

**Figure EV3.**
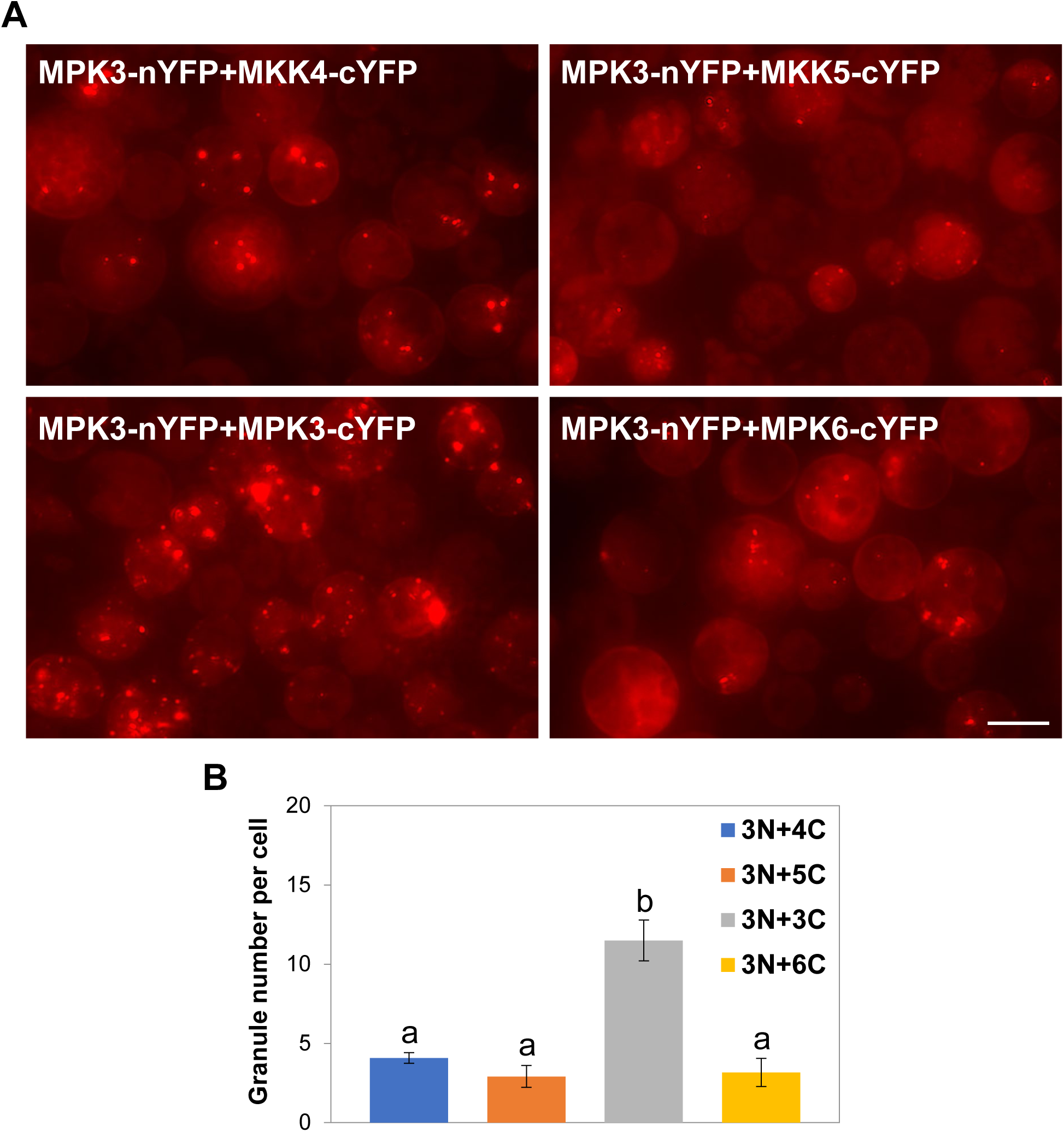
MPK3/6 and MKK4/5 interactions affect DCP1 granule assembly. **(A)** The granule assembly of co-expressed PB marker DCP1-mCherry is suppressed by hetero-dimers of MPK3-MKK4, MPK3-MKK5, and MPK3-MPK6 (right panel), whereas unaffected by homo-dimer of MPK3 (lower left panel), in BiFC analyses. Scale bar= 15 μm. Images of the BiFC signals (YFP) are not shown. **(B)** Quantitative analysis of granule number per cell as shown in (A). Columns represent means ± *SE*. Different letters above the bars indicate significant differences as indicated by ANOVA (*P* < 0.05).

**Figure EV4.**
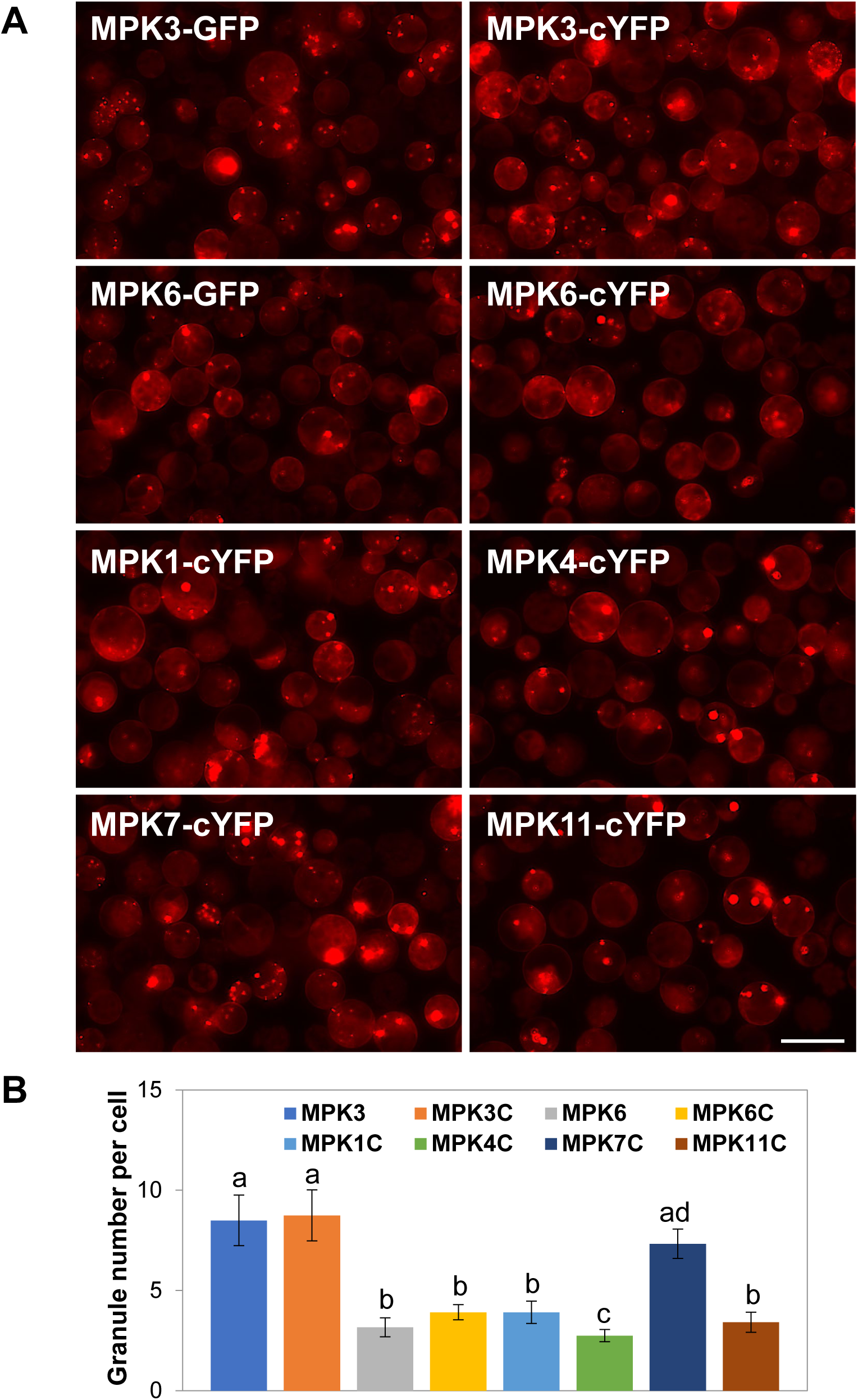
MPKs affect DCP1-mCherry granule dynamics. **(A)** Images of co-expressed GFP-MPK3/6 are not shown. MPKs-cYFP are single BIFC constructs not being able to generate yellow fluorescence signals. All the images were taken with the same exposure time. Scale bar= 25 μm. **(B)** Quantitative analysis of granule number per cell as shown in (A). Columns represent means ± *SE*. Different letters above the bars indicate significant differences as indicated by ANOVA (*P* < 0.05).

**Figure EV5.**
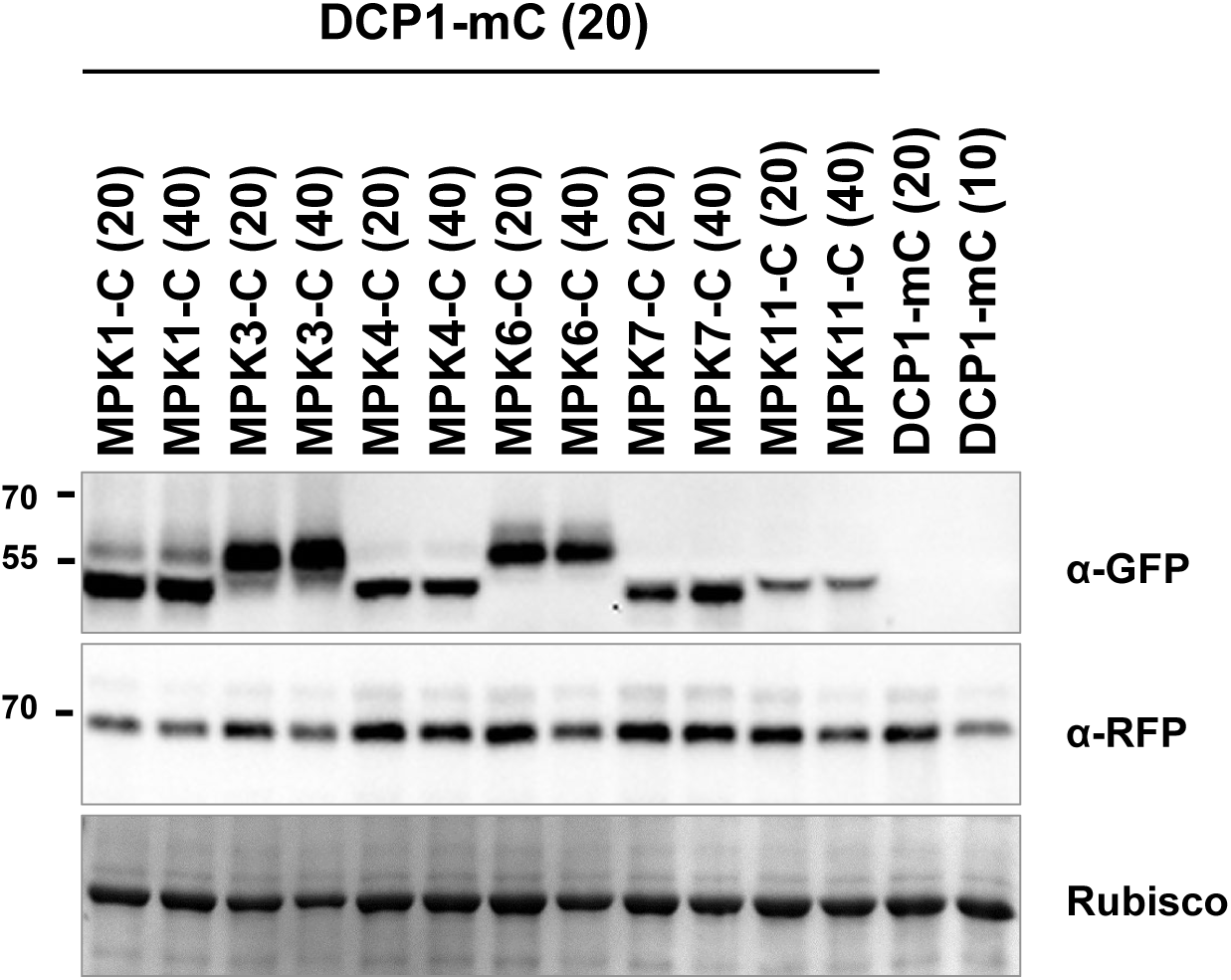
**DCP1-mCherry accumulation affected by MPKs and MKKs.** DCP1-mCherry was co-expressed with two different doses (20 vs 40 mg of plasmid) of MPK-cYFP in a protoplast transient expression assay. Immunoblot analysis was conducted using protein samples as indicated. GFP tagged proteins and MPK-cYFP were detected by GFP antibody and DCP1-mCherry was detected by RFP antibody.

**Figure EV6.**
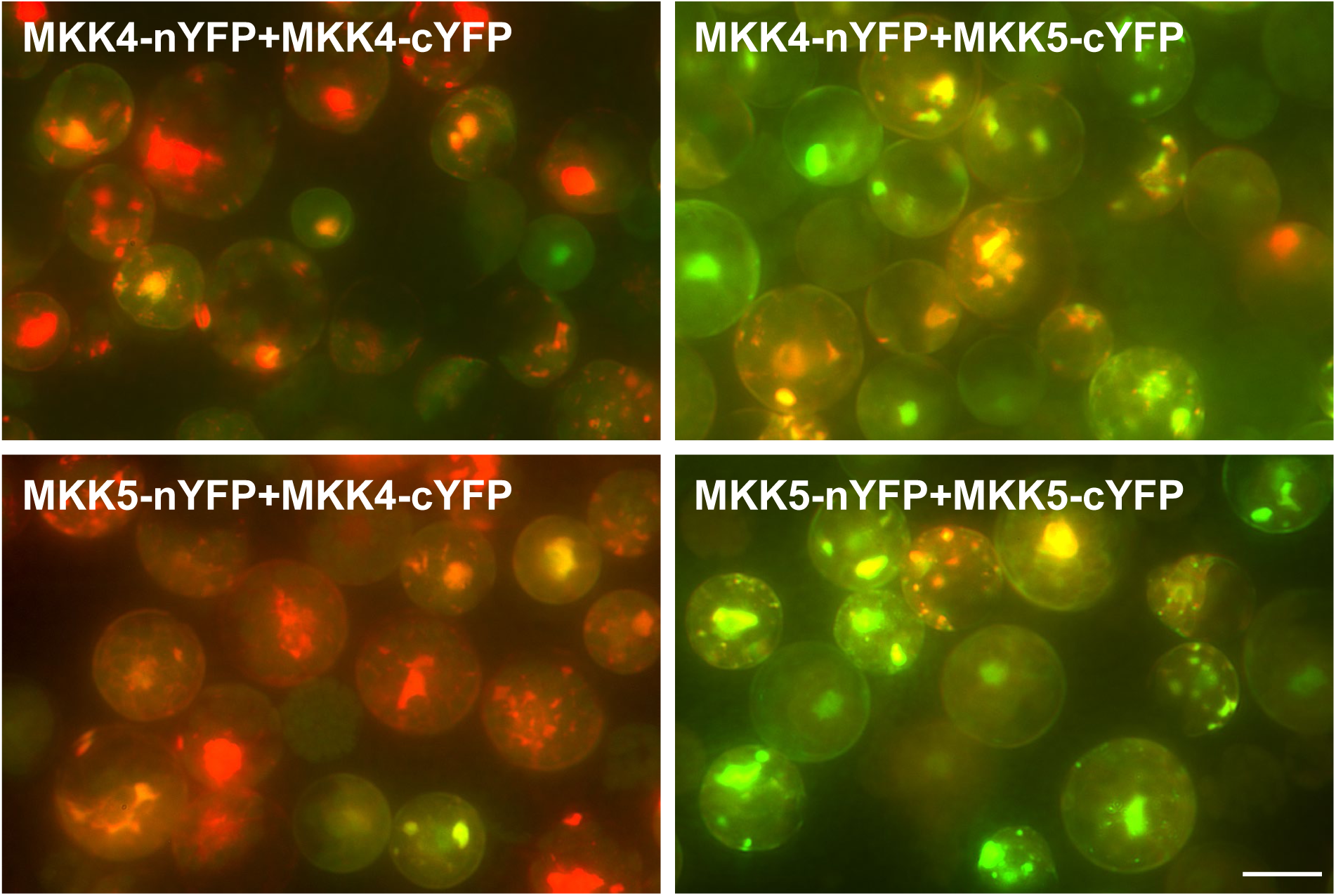
**As revealed by the dominant red signals, co-expression of SG marker UBP1b- mCherry suppresses the homo-dimerization of MKK4 (upper left) and hetero-dimerization of MKK5-MKK4 (lower left), but not the hetero-dimerization of MKK4-MKK5 (upper right) and homodimerization of MKK5 (lower right) in BiFC analyses.** Shown are merged images of BiFC (green signal from YFP) and SG marker (red signal from UBP1b- mCherry). Scale bar= 15 μm. Separate images from green and red channels are not shown.

**Figure EV7.**
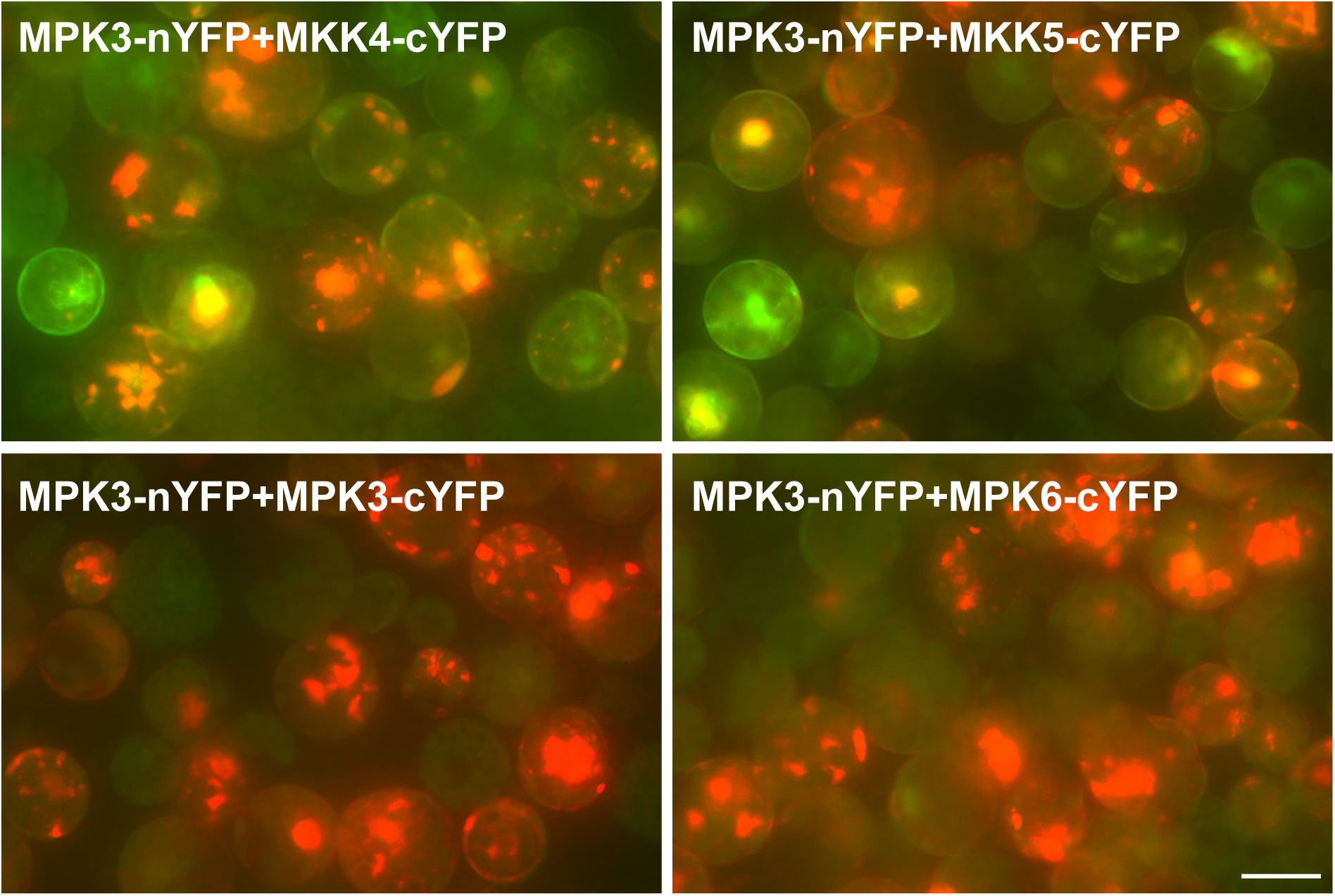
**As revealed by dominant red signals, co-expression of SG marker UBP1b-mCherry suppresses the homo-dimerization of MPK3 (lower left) and hetero-dimerization of MPK3-MPK6 (lower right), but to a lesser degree the hetero-dimerization of MPK3-MKK4 (upper left) and MPK3-MKK5 (upper right) in BiFC analyses.** Shown are merged images from BiFC (green signal from YFP) and SG marker (red signal from UBP1b-mCherry). Scale bar= 15 μm. Separate images from green and red channels are not shown.

**Figure EV8.**
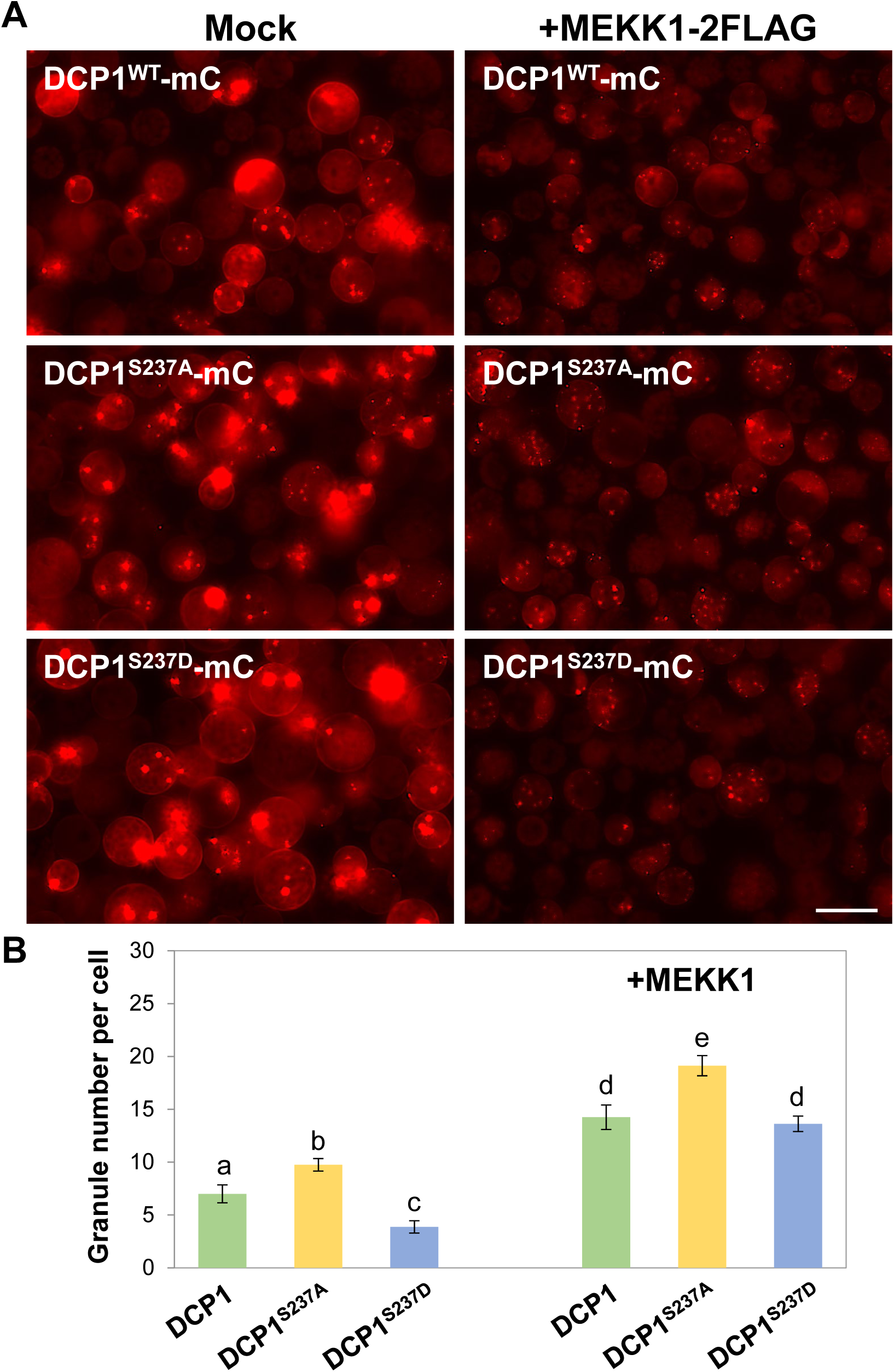
The MEEK1 enhances DCP1-mCherry granule assembly when co-expressing in an Arabidopsis protoplast transient expression analysis. **(A)** The enhancement is more pronunced for phospho-dead DCP1^S237A^ but not phospho-mimetic DCP1^S237D^ form. All the red images were taken with the same exposure time. Scale bar= 20 μm. **(B)** Quantitative analysis of typical small granule number per cell as shown in (A). Columns represent means ± *SE*. Different letters above the bars indicate significant differences as indicated by ANOVA (*P* < 0.05).

